# Barcoded reciprocal hemizygosity analysis via sequencing illuminates the complex genetic basis of yeast thermotolerance

**DOI:** 10.1101/2021.07.26.453780

**Authors:** Melanie B. Abrams, Julie N. Chuong, Faisal AlZaben, Claire A. Dubin, Jeffrey M. Skerker, Rachel B. Brem

## Abstract

Decades of successes in statistical genetics have revealed the molecular underpinnings of traits as they vary across individuals of a given species. But standard methods in the field can’t be applied to divergences between reproductively isolated taxa. Genome-wide reciprocal hemizygosity mapping (RH-seq), a mutagenesis screen in an inter-species hybrid background, holds promise as a method to accelerate the progress of interspecies genetics research. Here we describe an improvement to RH-seq in which mutants harbor barcodes for cheap and straightforward sequencing after selection in a condition of interest. As a proof of concept for the new tool, we carried out genetic dissection of the difference in thermotolerance between two reproductively isolated budding yeast species. Experimental screening identified dozens of candidate loci at which variation between the species contributed to the thermotolerance trait. Hits were enriched for mitosis genes and other housekeeping factors, and among them were multiple loci with robust sequence signatures of positive selection. Together, these results shed new light on the mechanisms by which evolution solved the problems of cell survival and division at high temperature in the yeast clade, and they illustrate the power of the barcoded RH-seq approach.

## INTRODUCTION

Understanding how and why organisms from the wild exhibit different traits is a central goal of modern genetics. Linkage and association mapping have driven decades of success in dissecting trait variation across individuals of a given species (Ott et al. 2015; Tam et al. 2019). But since these methods can’t be applied to reproductively isolated taxa, progress in the field of interspecies genetics has lagged behind. However, newer statistical-genetic methods appropriate to comparisons between species have been proposed in the recent literature (Weiss and Brem 2019), which hold promise for elucidating the genetics of ancient traits. For most such methods, limitations accruing from throughput and/or coverage issues remain to be refined.

The budding yeast *Saccharomyces cerevisiae* grows better at high temperature than any other species in its clade (Sweeney et al. 2004; Gonçalves et al. 2011; Salvadó et al. 2011; Hittinger 2013; Weiss et al. 2018), in keeping with its likely ecological origin in hot, East Asian locales (Peter et al. 2018). This derived and putatively adaptive trait serves as a model for the genetic study of deep evolutionary divergences. Thermosensitivity, the ancestral phenotype in the clade, is borne out in *S. paradoxus*, a close sister species to *S. cerevisiae*, making the former a useful point of comparison. Our group previously used this system as a testbed to develop RH-seq (Weiss et al. 2018), a genomic version of the reciprocal hemizygosity test (Stern 2014) that is well-suited to the mapping of natural trait variation between sister species. This technique starts with the generation of large numbers of random transposon mutant clones of a viable but sterile interspecies hybrid. In a given clone, loss of function from a transposon insertion in one species’ allele of a gene reveals the function of the uncovered allele from the other species.

These hemizygotes are competed en masse in a condition of interest; the abundance of each hemizygote in turn in the selected pool is quantified by bulk sequencing, and used in a test for allelic impact on the focal trait. In previous work, we identified eight genes through this approach at which species divergence contributed to thermotolerance (Weiss et al. 2018).

Against a backdrop of successful biological and evolutionary inference from our yeast RH-seq pilot (Weiss et al. 2018; Abrams et al. 2021), we noted that the combination of *S. cerevisiae* alleles of all eight genes mapped to thermotolerance recapitulated only <20% of the difference between the species (AlZaben et al. 2021). Thus, many of the determinants of yeast thermotolerance likely remain undetected. If so, boosting the replication and throughput of genetic mapping, to enable higher statistical power, could help meet the challenge. In our initial implementation of RH-seq, we had quantified the abundance of hemizygotes in a sample by sequencing across the transposon junction with the genome, using one universal primer that recognized the transposon and another recognizing a ligated adapter at DNA fragment ends (Weiss et al. 2018). This protocol, though rigorous, is labor-intensive and expensive, limiting the potential for throughput and coverage. A higher-throughput alternative starts with the tagging of transposon sequences by random short DNA barcodes (Wetmore et al. 2015). After mutagenesis of a genotype of interest by these barcoded transposons, and then selection of the mutants in bulk in a challenging condition, mutant abundance can be quantified from sequencing of DNA straight from the pool with a simple PCR. We set out to adapt this barcoding strategy to enable highly replicated RH-seq, with application to yeast thermotolerance as a test case to achieve a deeper exploration of the complex genetics of the trait.

## MATERIALS AND METHODS

### Construction of a randomly barcoded piggyBac transposase pool

For barcoded RH-seq, we constructed a pool of plasmids, each harboring the piggyBac transposase and a randomly barcoded copy of the piggyBac transposon, via Golden Gate cloning of random 20bp barcodes flanked by universal priming sites into a plasmid backbone containing the piggyBac machinery, modified from pJR487 (Weiss et al. 2018) as follows (Figure S1).

#### Preparation of the backbone vector

To allow the use of BbsI as the Type IIS restriction enzyme for Golden Gate cloning of barcodes into pJR487 (see below), we first removed all three BbsI cut sites from pJR487 by introducing silent mutations that disrupted the restriction enzyme’s recognition pattern. The resulting plasmid was called pCW328. We next modified pCW328 to make a Golden-Gate-ready vector, with the final identifier pJC31, by replacing transposon nucleotides with those of a stuffer at a location 70 nucleotides from the end of the right arm of the transposon (Table S1); see Supplementary Note and Figure S2 for a description of this choice. The stuffer contained two BbsI cut sites with custom Type IIS overhang sequences from (Lee et al. 2015), and a NotI cut site in between the two BbsI cut sites. All cloning steps were carried out by GenScript, Inc.

**Table 1.**
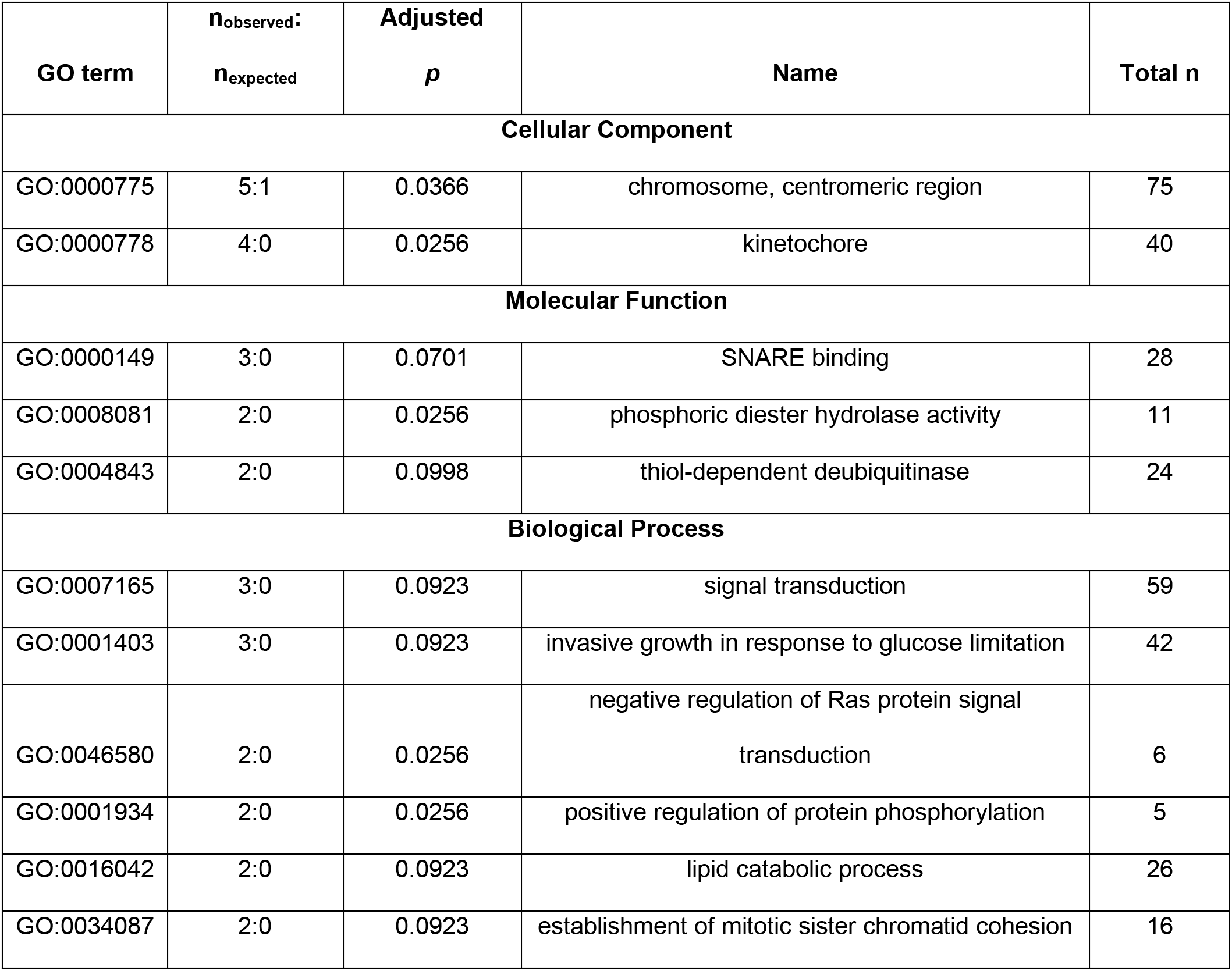
Functional enrichment among thermotolerance loci. Each row with numerical data reports a Gene Ontology (GO) term enriched for RH-seq hit genes. n_observed_, the number of genes from among top hits from thermotolerance RH-seq that were annotated with the term. n_expected_, the number of genes annotated with the term in the same number of randomly chosen genes from the genome, as a median across samples. Adjusted *p*, resampling-based significance of the enrichment after Benjamini-Hochberg correction. Total *n*, the total number of genes annotated in the GO term in *S. cerevisiae*.

| GO term | $n_{\text{observed}}$<br>$n_{\text{expected}}$ | Adjusted<br>$p$ | Name | Total $n$ |
| --- | --- | --- | --- | --- |
| Cellular Component |  |  |  |  |
| GO:0000775 | 5:1 | 0.0366 | chromosome, centromeric region | 75 |
| GO:0000778 | 4:0 | 0.0256 | kinetochore | 40 |
| Molecular Function |  |  |  |  |
| GO:0000149 | 3:0 | 0.0701 | SNARE binding | 28 |
| GO:0008081 | 2:0 | 0.0256 | phosphoric diester hydrolase activity | 11 |
| GO:0004843 | 2:0 | 0.0998 | thiol-dependent deubiquitinase | 24 |
| Biological Process |  |  |  |  |
| GO:0007165 | 3:0 | 0.0923 | signal transduction | 59 |
| GO:0001403 | 3:0 | 0.0923 | invasive growth in response to glucose limitation | 42 |
| GO:0046580 | 2:0 | 0.0256 | negative regulation of Ras protein signal<br>transduction | 6 |
| GO:0001934 | 2:0 | 0.0256 | positive regulation of protein phosphorylation | 5 |
| GO:0016042 | 2:0 | 0.0923 | lipid catabolic process | 26 |
| GO:0034087 | 2:0 | 0.0923 | establishment of mitotic sister chromatid cohesion | 16 |

#### Preparation of barcode oligonucleotides

To make barcodes, we acquired an oligonucleotide pool from IDT that contained random 20 bp sequences (from hand-mixed random nucleotides) flanked by universal priming regions, U1 and U2 (Wetmore et al. 2015, Coradetti et al. 2018). These custom oligos were produced and PAGE purified by IDT. Additionally, we designed forward (FW_BbsI_JC) and reverse (REV_BbsI_JC) primers which each contained a BbsI cut site, BbsI overhang sequences complementary to the backbone vector, and either universal priming sequence (Table S2) (Coradetti et al. 2018). We set up 50 μL amplification PCR reactions with 1 μL of random 20 bp barcodes as template, from a 2.5 μM stock, and 0.25 μL of each of the forward and reverse primers from a 100 μM stock.

Amplification used Phusion High Fidelity polymerase (NEB) and the following cycling protocol: 98°C for 30 seconds, (98°C for 10 seconds, 58°C for 30 seconds, 72°C for 60 seconds) × 6, 72°C for five minutes. PCR products were purified (Zymo DNA Clean & Concentrator kit) and then combined. This yielded the final donor barcodes: random 20bp barcodes flanked by universal priming regions, with BbsI cut sites at the extreme edges.

#### Cloning barcodes into plasmids

To clone barcodes into pJC31, we proceeded in two barcoding reactions. The first reaction contained 2:1 molar ratio of vector to barcodes (4 μg of pJC31 and 128 ng of donor barcodes), 5 μL of 10X T4 Ligase Buffer (ThermoFisher), 2.5 μL of T4 Ligase (ThermoFisher), 2.5 μL FastDigest Bpil (ThermoFisher), and sterile water up to 50 µL. The cycling program was: 37°C for five minutes, (37°C for two minutes, 16°C for five minutes) x 25, 65°C for 10 minutes. Then a mixture containing 5 μL 10X FastDigest Buffer (ThermoFisher), 3.13 μL BSA 2 mg/mL (NEB), 12.5 μL FastDigest NotI (ThermoFisher), and 12.5 μL FastDigest Bpil (ThermoFisher) was spiked into the reaction and incubated at 37°C for 16 hours to digest unbarcoded backbone vectors. Ten of these reactions were combined, purified, and eluted in H_2_O (Zymo DNA Clean & Concentrator). To spot-check this cloning, 5 μL of this product was transformed into 25 μL of *E. coli* 10beta electrocompetent cells (NEB). Sanger sequences across the barcode regions of 20 individually miniprepped *E. coli* colonies showed 95% barcoding efficiency.

The second reaction contained 2:1 molar ratio of vector to donor barcodes (4μg of pJC31 and 128 ng of donor barcodes), 5 μL of 10X T4 Buffer (ThermoFisher), 2.5 μL T4 Ligase (ThermoFisher), 2.5 μL Bpil (ThermoFisher), and sterile water up to 50 µL. The cycling program was: 37°C for five minutes, (37°C for two minutes, 16°C for five minutes) x 25, 65°C for 10 minutes. Then a mixture containing 2.5 μL 10X FastDigest Buffer (ThermoFisher), 2.5 μL G Buffer, (ThermoFisher), 3.13 μL BSA 2 mg/mL (NEB), 12.5 μL FastDigest NotI (ThermoFisher), and 12.5 μL Bpil (ThermoFisher) was spiked in the reaction and incubated at 37°C for 16 hours to digest remaining unbarcoded backbone vectors. Six of these reactions were combined, purified, and eluted in H_2_O (Zymo DNA Clean & Concentrator). Then every 5 μL of cleaned eluted product was redigested with 5μL of NotI-HF (NEB), 5 μL 10X CutSmart buffer (NEB), and 35 μL H_2_O at 37°C for 16 hours then 80°C for 20 minutes. The reactions were purified again (Zymo DNA Clean & Concentrator) and pooled. Spot checks of this cloning reaction proceeded as above, and Sanger sequences across the barcode regions of 20 individually miniprepped *E. coli* colonies showed 95% barcoding efficiency.

Purified plasmids from the two reactions were combined in a master tube of DNA before transforming into electrocompetent *E. coli* cells (NEB) to generate the final barcoded piggyBac pool (final identifier P58). Each electroporation cuvette (BTX) contained 25 μL of 10beta electrocompetent cells (NEB) and 5 μL of cleaned master tube DNA from the previous golden gate barcoding step. We performed 21 electroporation reactions in total using the Bio-Rad GenePulser Xcell machine set to 2.0 kV, 200 Ohms, 25 μF. After electroporation, each culture was recovered in provided outgrowth media (NEB) by shaking at 37°C at 250 rpm for 1.5 hours. After recovery, all independent 21 electroporation reactions were combined.

The combined recovered transformation *E. coli* culture was used to inoculate two 1L fresh LB cultures containing carbenicillin at 100 μg/mL to select for *E. coli* cells containing barcoded piggyBac plasmids. Each culture was incubated for 15.5 hours at 37°C, 250 rpm (overnight) to expand the barcoded piggyBac *E. coli* pool. Then the two cultures were combined yielding the final barcoded transposon plasmid pool, P58. This was aliquoted into 1 mL volumes with 15% glycerol and stored at -80°C.

#### Sequencing verification of barcoded piggyBac pool plasmid DNA for barcode diversity

To verify barcode diversity in the barcoded piggyBac plasmid pool (P58), we sequenced barcodes as follows. One frozen aliquot of P58 was inoculated into 1.25 L of LB containing carbenicillin 100 μg/mL and grown for 16 hours 37°C, 250 rpm or until it reached an OD_600_ of 2.1. This culture was gigaprepped on using a column kit (Invitrogen) to generate 5 mg of plasmid. We used this as input into a PCR with primers (Table S2) annealing to the universal priming regions flanking the barcode. These primers were dual-indexed, although in this work we only carried out sequencing of the resulting amplicon from one end (see below), such that only one index was used. The generic form of the forward primer was AATGATACGGCGACCACCGAGATCTACACTCTTTCCCTACACGACGCTCTTCCGATCT(N1- 4)xxxxxxGTCGACCTGCAGCGTACG, where the N1-4 represent variable amounts of random bases from 1-4 to help samples cluster on the Illumina lane and the (x6) represent a unique 6- bp index sequence for multiplexing samples. The generic reverse primer was CAAGCAGAAGACGGCATACGAGATxxxxxxGTGACTGGAGTTCAGACGTGTGCTCTTCCGAT CTGATGTCCACGAGGTCTCT . Four PCR reactions used 50 ng of prepped P58 plasmid template each. Amplification used Q5 High Fidelity Polymerase (NEB) and a cycling program 98°C for four minutes, (98°C for 30 seconds, 55°C for 30 seconds, 72°C for 30 seconds) x 25, 72°C for five minutes. Each PCR product was purified on a column (Zymo DNA Clean & Concentrator-5 Kit) and eluted in 10 μL prewarmed 65°C provided elution buffer (Zymo). Six μL of each were then combined and sequenced off the U2 region via Illumina amplicon sequencing, on one lane of HiSeq4000 SR50 at the Genomics Sequencing Laboratory at UC Berkeley. Reads sequenced per library are reported in Table S3. Sequencing of the *E. coli* vector pool p58 revealed 27,538,142 barcodes with an estimated sequencing error rate of 1.38% analyzed as described (Coradetti et al. 2018).

### Yeast hemizygote pool construction via barcoded transposon mutagenesis

We constructed our yeast hemizygote pool essentially as described (Weiss et al. 2018) but with modifications as follows.

To prepare plasmid DNA for mutagenesis, one frozen aliquot of P58 was inoculated into 1.25L of LB containing carbenicillin 100 μg ml^−1^ and grown for 16 hours at 37°C, 250 rpm or until it reached an OD_600_/mL of 2.1. This culture was gigaprepped on using a column kit (Invitrogen) to generate 5 mg of plasmid.

Next, we transformed yeast in several, smaller subpools which we combined to form a final pool as follows. We carried out mutagenesis of CW27, an F1 hybrid from the mating of *S. cerevisiae* DBVPG1373 with *S. paradoxus* Z1 (Weiss et al. 2018) across the first two days. The first day, we generated one subpool in a single 50 mL culture and one subpool in five 50 mL cultures at OD_600_/mL ∼0.9 (∼45 OD_600_ units of cells each). The second day, we generated two subpools in five 50 mL cultures each at OD_600_/mL ∼0.9 (∼45 OD_600_ units of cells).

To generate subpools consisting of a single 50 mL culture, one colony of CW27 was inoculated into 5 mL of YPD and incubated at 28°C 200 rpm. 24 hours later, the OD_600_/mL of the overnight culture was 3.86. It was backdiluted to an OD_600_/mL of 0.1 in 50 mL of YPD in a 250 mL Erlenmeyer flask and grown with shaking at 28°C, 200 rpm for 5.5 hours. After 5.5 hours, it had reached OD_600_/mL ∼0.9 and cells were at mid-log phase. This 50 mL culture was gently pelleted at 1000xg for three minutes. The pellet was washed with 25 mL sterile water and then 5 mL of 0.1 M lithium acetate (Sigma) mixed with 1X Tris-EDTA buffer (10 mM Tris-HCl and 1.0 mM EDTA); after spin-down, to the tube was added a solution of 0.269 mg of P58 mixed 5:1 by volume with salmon sperm DNA (Invitrogen), followed by 3 mL of 39.52% polyethylene glycol, 0.12 M lithium acetate and 1.2X Tris-EDTA buffer (12 mM Tris-HCl and 1.2 mM EDTA). The tube was rested for 10 minutes at room temperature, then heat-shocked in a water bath at 37°C for 26 minutes. The tube was gently spun at 1000g for three minutes after which supernatant was removed. We transferred the cells to a flask and added YPD to attain an OD_600_/mL of ∼0.35–4 in ∼70 mL. Each such culture was recovered by shaking at 28°C and 200 rpm for two hours. G418 (Geneticin; Gibco) was added to each at a concentration of 300 μg/mL to select for those cells that had taken up the plasmid, and cultures were incubated with 200 rpm shaking at 28°C for two days until each reached an OD_600_/mL of ∼2.5. We transferred cells from this culture, and YPD + G418 (300 μg/mL), to new 250 mL flasks at the volumes required to attain an OD_600_/mL of 0.2 in 50 mL each. We cultured each flask with 200 rpm. shaking at 28°C overnight until each reached an OD_600_/mL of 3.43. To cure transformants of the P58 URA3+ plasmid, we spun down 10% of this master culture, and resuspended in water with the volume required to attain a cell density of 1.85 OD_600_/mL. Four mL of this resuspension were plated (1 mL per 24.1 cm x 24.1 cm plate) onto plates containing complete synthetic media with 5- fluorooritic acid (0.2% dropout amino acid mix without uracil or yeast nitrogen base (US Biological), 0.005% uracil (Sigma), 2% D-glucose (Sigma), 0.67% yeast nitrogen base without amino acids (Difco), 0.075% 5-fluorooritic acid (Zymo Research)). After incubation at 28°C to enable colony growth, colonies were scraped off all four plates and combined into water at the volume required to attain 44 OD_600_/mL, yielding the transposon mutant hemizygote subpool. This was aliquoted into 1 mL volumes with 10% dimethylsulfoxide and frozen at −80°C.

To generate subpools consisting of five 50 mL cultures, one colony of CW27 was inoculated to 100 mL of YPD in a 250 mL Erlenmeyer flask and incubated shaking at 28°C, 200 rpm. Twenty-four hours later, the OD_600_/mL of the overnight culture was OD_600_/mL 3.89. The overnight culture was backdiluted to OD_600_/mL 0.1 in 250 mL of YPD and incubated for 5.5 hours at 28°C, 200 rpm. After 5.5 hours, the OD_600_/mL reached 0.9 and cells were split into five 50 mL conical tubes, and subjected each to heat shock as above. We then transferred all cells from this post-transformation culture to one 1L flask and added fresh YPD to attain OD_600_/mL 0.4 in ∼750 mL YPD. The transformed culture was recovered by shaking at 28°C, 200rpm, for two hours. G418 (300mg/ul) was added to select for the transposed cells. The culture continued shaking for 48 hours or until the OD_600_/mL reached 2.1. This culture was then backdiluted to create a new culture at OD_600_/mL 0.2 in 500 mL of YPD with 300mg/μL G418 shaking for 24 hours at 28°C, 200 rpm until it reached OD_600_/mL ∼3.4 The curing, scraping, and freezing steps were the same as above.

To combine the four subpools to yield the final 160X hemizygote pool (final identifier P75), three 1 mL aliquots of each subpool were thawed on ice for one hour. They were transferred to each of four 1L flasks with 500 mL YPD to OD_600_/mL 0.2, cultured at 28°C, 200 rpm for 17 hours upon which the OD_600_/mL was 3.5-4. They were gently pelleted, combined, and resuspended in two ways to reach OD_600_/mL of 44: YPD with 15% glycerol and YPD with 7% DMSO, aliquoted to 1 mL volumes, and frozen at -80°C.

### Tn-seq mapping of yeast hemizygote pool

#### Tn-seq library preparation

To associate barcoded transposon insertions to genomic location in the hemizygote pool, which we refer to as Tn-seq, we first sequenced barcoded transposon insertions according to the methods of (Weiss et al. 2018) as follows. Each 44 OD_600_/mL aliquot of each subpool or final pool was thawed on ice, and its genomic DNA (gDNA) was harvested with the ZR Fungal/Bacterial DNA MiniPrep Kit (Zymo Research). gDNA was resuspended in DNA elution buffer (Zymo Research) prewarmed to 65°C, and its concentration was quantified using a Qubit 4.0 fluorometer. Illumina transposon sequencing (Tn-seq) library construction was as described previously. Briefly, gDNA was sonicated and ligated with common adapters, and for each fragment deriving from a barcoded transposon insertion in the genome, a sequence containing a barcode, a portion of the transposon, and a portion of its genomic context (the barcoded transposon–genome junction) was amplified using one primer homologous to the U1 region immediately upstream of barcode and another primer homologous to a region in the adapter. See Table S2 for the transposon-specific primer (“forward primer”), where Ns represent random nucleotides, and the indexed adapter-specific primer (“reverse primer”). Amplification used Jumpstart polymerase (Sigma) and the following cycling protocol: 94°C for two minutes, (94°C for 30 seconds, 65°C for 20 seconds, 72°C for 30 seconds) × 25, 72°C for 10 minutes.

Sequencing of paired-end reads of 150 bp was done over two lanes on a HiSeq4000 at Novogene Corporation (Sacramento, CA) and one lane on a NovaSeq SP at the Genomics Sequencing Laboratory at UC Berkeley (Berkeley, CA). Reads sequenced per library are reported in Table S4.

#### Tn-seq data analysis

Tn-seq data of the hemizygote pool was analyzed, to infer transposon insertions on the basis of barcodes detected in reads as junctions with genomic sequence, essentially as described (Coradetti et al. 2018) (https://github.com/stcoradetti/RBseq/tree/master/Old_Versions/1.1.4), with the following modifications. For each barcode, instead of scanning positions for the end of the insertion from a sequence specified by a model file, we searched for the final 22 base pairs of the right arm of the piggyBac transposon allowing for two mismatches. For annotation, we converted the annotation file from https://github.com/weiss19/rh-seq for the *S.* cerevisiae D1373 x *S. paradoxus* Z1 hybrid to a compliant GFF3 file using Another GFF Analysis Toolkit (AGAT) - Version: v0.4.0 (https://github.com/NBISweden/AGAT). Then, we used a custom Jupyter notebook to annotate the file generated by the RBseq mapping software.

Quality control for Tn-seq, to eliminate barcodes whose junction genomic sequence mapped to multiple insertion locations in the hybrid genome, and to minimize the proportion of sequencing errors included in final tallies, was as described (Coradetti et al. 2018). Briefly, we eliminated from further consideration any case where a barcode observed in Tn-seq sequencing data differed from another, much more abundant, barcode by a single base (a total of 2,024,812 off-by-one barcodes in 2,888,129 reads). We also filtered out off-by-two barcodes (280,949 barcodes in total). Separately, we eliminated barcodes that were detected in sequencing data as a junction with more than one genomic context, suggesting the respective transposon had inserted into multiple locations in one or many clones (98,669 barcodes where this inference was based on multiple strong mapping matches, and an additional 46,583 barcodes where this inference was ambiguous, with one strong mapping match with reads outnumbered by those assigned to weaker mapping matches). The final filtered barcode set comprised 548,129 uniquely barcoded and mapped inferred transposon insertions in the P75 hemizygote pool, at an average read depth of 308.6 reads, and a median read depth of 47 reads; 166,834 of these insertions were mapped as genic. The annotation script, GFF3 file, and modified mapping script are available at https://github.com/melanieabrams-pub/RH-seq_with_barcoding.

### Competition cultures

For the thermotolerance competition at 37°C (Table S5), one aliquot of the yeast hemizygote pool was thawed and inoculated into 150 mL of YPD in a 250 mL unbaffled Erlenmeyer flask and grown for six hours at 28°C, 200 rpm. This pre-culture (T_0_, at OD_600_/mL of 1.22) was backdiluted into 12 10 mL competition cultures at 200 rpm at each of 28°C and 37°C, with a starting OD_600_/mL of 0.02 or 0.05 in at 28°C and 37°C respectively. These competition cultures were maintained within logarithmic growth through back-dilutions into fresh tubes of 10 mL of YPD at the same optical density as the starting culture, for a total of 10-15 generations. Dilutions for the 28°C competition cultures were performed after 8.5, 18.5, and 25.5 hours after the T_0_ timepoint, and dilutions for the 37°C competition cultures were performed after 8.5, 18.5, 25.5 hours and 32.5 hours after the T_0_ timepoint. The entire cell culture was harvested from each of these biological replicate tubes for sequencing as biological replicates.

For thermotolerance competition at 36°C (Table S6), competition cultures were grown as above with the following differences. The high temperature was 36°C, instead of 37°C. The pre-culture (T_0_, at OD_600_/mL of 0.693 after 5.5 hours at 28°C, 200 rpm) was backdiluted to a starting OD_600_/mL of 0.02 for competition cultures at 36°C. Dilutions for both the 28°C and 36°C competition cultures were performed after 8.5, 18.5 and 25.25 hours after the T_0_ timepoint.

Eleven instead of 12 replicates were carried out at 28°C.

### Barcode quantification from competition cultures

#### Bar-seq library preparation

To determine the abundance of barcoded transposon mutant hemizygote clones after selection, we sequenced barcodes insertions as follows. Each cell pellet from a selection sample was thawed on ice, and its genomic DNA (gDNA) was harvested with the Zymo QuickDNA Kit (Zymo#D6005). gDNA was resuspending in DNA elution buffer (Zymo Research) prewarmed to 65°C, and its concentration was quantified using a Qubit 4.0 fluorometer. The barcode insertion was amplified as above (see *Sequencing verification of barcoded piggyBac pool plasmid DNA for barcode diversity*). Each PCR product was purified on a column (Zymo DNA Clean & Concentrator) and eluted in 10 μL prewarmed 65°C provided elution buffer (Zymo). Six μL of each were then combined and sequenced off the U2 region by Illumina sequencing on one lane of HiSeq4000 SR50 at the QB3 Genomics Sequencing Laboratory at UC Berkeley.

#### Bar-seq data analysis

Bar-seq mapping and quantification were as described (Coradetti et al. 2018) (https://github.com/stcoradetti/RBseq/tree/master/Old_Versions/1.1.4), wherein only barcodes that passed quality control in Tn-seq (see *Tn-seq data analysis* above) were analyzed for quantitative measures of abundance via Bar-seq. Thus we did not use in our screen any barcode that was detected in Bar-seq sequence data but not Tn-seq data (the product of *e.g.* sequencing errors in Bar-seq, or a failure to observe in Tn-seq a barcode associated with a bona fide transposon insertion that could be detected in Bar-seq). A total of 301,349 barcodes conformed to these criteria from across all replicates of Bar-seq in competitions for the dissection of determinants of growth at 37°C relative to 28°C, with an average read depth of 305.3 reads and a median of 12 reads; 89,772 of these Bar-seq detected barcodes corresponded to inferred transposon insertions in genes and were analyzed as input to the reciprocal hemizygosity testing pipeline described below. In a given replicate competition culture we detected a median 1 x 10^5^ barcodes. The latter represented a fifth of the size of the total pool of hemizygotes detectable after quality control by Tn-seq (5.5 x 10^5^; see *Tn-seq data analysis* above). Thus, the extent of bottlenecking in any given competition experiment was modest, with diversity retained at the order of magnitude of the mutant pool size.

Competitions for the dissection of growth at 36°C relative to 28°C (Table S6) used the same procedures as above, mapping a total of 230,469 barcodes, 68,523 of which corresponded to inserts in genes and were analyzed as input to the reciprocal hemizygosity testing pipeline described below. In a given replicate competition culture, we detected a median 5 x 10^4^ barcodes.

### Reciprocal hemizygosity testing

The tabulated counts of abundance from Bar-seq for each barcode in each replicate were used as input into reciprocal hemizygosity tests essentially as in (Abrams et al. 2021), with slight changes as follows. We had in hand each barcode which had been sequenced as a junction with a unique genomic location in the Tn-seq step and had passed quality control there (see *Tn-seq data analysis* above), and which was now detected in competition cultures. We interpreted each such barcode as reporting a hemizygote clone bearing a transposon insertion at the respective position of the respective species’ allele (*S. cerevisiae* or *S. paradoxus*), with the other species’ allele retained as wild-type at that locus. In what follows, we refer to each such barcode as reporting an inferred hemizygote clone, with respect to its growth behavior in competition cultures. As in (Abrams et al. 2021), for a given biological replicate we normalized the abundances attributed to each inferred hemizygote genotype to the total number of sequencing reads in the respective sample, and we eliminated from further analysis insertions which had been annotated as intergenic, or as corresponding to the plasmid used to generate this library. For reciprocal hemizygosity tests, we excluded from consideration any gene with fewer than three inferred hemizygote genotypes per allele. Of the retained genes, for each inferred hemizygote genotype, we tabulated the quantity a_experimental,i_, the sequencing-based abundance measured after the competition culture in biological replicate *i* of growth at the experimental temperature (36 or 37°C), and, separately, we calculated a_control,i_, the analogous quantity from growth at the control temperature (28°C), for *i* = [1,12]. We then took the mean of the latter and used it to tabulate the temperature effect on the inferred hemizygote genotype in replicate *i*, *t_i_* = log_2_(a_experimental,i_/a_control,mean_). As in (Abrams et al. 2021), we eliminated an inferred hemizygote genotype if the coefficient of variation of this quantity exceeded 2.0, or there were fewer than 1.1 normalized reads. With the data for the remaining inferred hemizygote genotypes (Tables S5 and S6), for a given gene, we compiled the vector of the *t* measurements across replicates and all inferred hemizygote genotypes with each species’ allele of the hybrid disrupted in turn, and discarded genes where the coefficient of variation of the *t* measurements across hemizygote inserts for one or both alleles exceeded 10. For the remainder, we used the Mann-Whitney test to compare these two vectors, with Benjamini-Hochberg correction for multiple testing (Tables S7 and S8). For a given gene, we calculated the effect size as the difference between two values: the log_2_(abundance at the experimental temperature/abundance at 28°C) of the average inferred hemizygote genotype representing a transposon insertion in the *S. cerevisiae* allele, and the analogous quantity among inferred hemizygote genotypes representing insertions in the *S. paradoxus* allele of the gene. Scripts for this modified RH-seq analysis pipeline are available at https://github.com/melanieabrams-pub/RH-seq_with_barcoding. We earmarked top candidate genes for factors contributing to the thermotolerance of *S. cerevisiae* as those with corrected Mann-Whitney *p* < 0.05 in the reciprocal hemizygosity test, and an effect size < -0.5, *i.e.* disrupting the *S. cerevisiae* allele was associated with a strong defect in thermotolerance relative to disruption of the *S. paradoxus* allele; we refer to this gene set as our top barcoded RH-seq hit gene list.

### Analysis of inferred interactions between top hit genes from barcoded RH-seq

For the circos plot reporting inferred interactions between top hit genes from barcoded RH-seq, we used the STRING database (Szklarczyk et al. 2021), accessed September 30, 2021, which incorporates experimental/biochemical data from DIP, BioGRID, HPRD, IntAct, MINT, and PDB, and curated data from Biocarta, BioCyc, Gene Ontology, KEGG, and Reactome. Widths of edges between nodes in the circos plot represent STRING confidence scores, each the probability of a true positive interaction between a given two genes (Szklarczyk et al. 2021).

To test the encoded proteins of top barcoded RH-seq hit genes for enrichment of physical interactions with each other, we used curated known interactions from BioGRID (Oughtred et al. 2021) as housed in the Saccharomyces Gene Database, downloaded February 19, 2021. We tabulated the number of physical interactions between the proteins encoded by RH-seq hit genes, and we divided that by the total number of interactions involving one RH-seq hit gene and any other gene in the genome; call this ratio r_true_. Then, we drew a random sample of genes from the genome, as described above for GO term resampling. We tabulated, in this random gene set, the number of physical interactions between genes in that sample, and we divided that by the total number of interactions involving one gene in the random sample and any other gene in the genome, to yield r_resample_. We repeated this procedure 10,000 times, and we used the proportion of resampled groups where r_resample_ was greater than or equal to r_true_ as a one-sided *p* value assessing the significance of enrichment of interactions between our genes of interest.

### Gene Ontology analyses of top hit genes from barcoded RH-seq

To test top barcoded RH-seq hit genes for enrichment for overrepresentation of a particular Gene Ontology (GO) term, we mapped each gene to its Gene Ontology groups based on data from geneontology.com (Ashburner et al. 2000). We filtered out GO terms with fewer than five or with more than 200 gene members. We also filtered out GO terms with identical membership in the genome. We took the subset of the remaining GO terms with at least one member among our top barcoded RH-seq hit genes. Then, we randomly sampled genes from the genome, ensuring the same proportion of essential genes as in our set of top barcoded RH-seq hit genes based on the essentiality annotations of (Winzeler 1999). We tabulated whether our random sample had greater or fewer genes with that GO term than our candidate set. We repeated this procedure 10,000 times and used the proportion of these resampled groups that had more genes in the given GO term as the initial *p* value assessing the significance of the enrichment of that GO term. Then, we applied Benjamini-Hochberg correction for multiple hypothesis testing to generate final, adjusted *p*-values for the enrichment of the given GO term among top barcoded RH-seq hit genes.

To test Biological Process ontologies for enrichment for large magnitudes of the effect of allelic variation on thermotolerance, we used the latter as tabulated in **Reciprocal hemizygosity testing** above. We filtered GO terms as above, and then excluded all genes absent in our barcoded RH seq analysis, which would have no associated quantity for the effect of allelic variation to resample. For each retained term in turn, we first tabulated the median absolute value of the effect size of the gene members for which we had data, e_true_. Then, we tabulated the analogous quantity for a random sample of the same number of genes from the genome, e_resample_, ensuring the same proportion of essential genes as above. We repeated this procedure 100 times, and used the proportion of the resampled groups for which e_resample_ was greater than or equal to e_true_ as an initial *p* value assessing the enrichment of large effects of allelic variation in the genes the term. For all GO terms with an initial *p* value < 0.1, we repeated this procedure 10,000 times to calculate a more precise *p* value. Then, we applied the Benjamini-Hochberg correction for multiple hypothesis testing to generate final, adjusted *p*-values for the enrichment of the given GO term for large effects of allelic variation on thermotolerance.

### Molecular evolution analysis of RH-seq hit genes

Branch length PAML analysis with codeML was performed as in (Dubin et al. 2020). Hits were manually inspected for the quality of the alignment, and one, *YAL026C*, was discarded for poor alignment quality leading to an artifactually high branch length. We used the inferred branch lengths as input into a resampling test as in **Gene Ontology analyses of top hit genes from barcoded RH-seq** above, and we performed a one-sided significance test for long branch lengths along the *S. cerevisiae* lineage. Branch-site PAML analysis with codeML was performed as in (Abrams et al. 2021). Jalview version 2 was used to visualize the percent identity of amino acid sequence alignments (Waterhouse et al. 2009). McDonald-Kreitman analysis statistics were calculated as in (Abrams et al. 2021). Fisher’s exact test was used to compute *p*-values for individual loci, and these were adjusted using the Benjamini-Hochberg correction for multiple hypothesis testing.

### Data availability statement

Sequencing data are deposited in the Sequence Read Archive under the accession PRJNA735401. Strains and plasmids are available upon request. Custom scripts for the barcoded RH-seq analysis are available at https://github.com/melanieabrams-pub/RH-seq_with_barcoding. The authors affirm that all data necessary for confirming the conclusions of the article are present within the article, figures, and tables.

## RESULTS

### Dissecting thermotolerance divergence between species by barcoded transposon mutagenesis

With the goals of boosting RH-seq throughput and power, and achieving new insights into the genetics and evolution of yeast thermotolerance, we set out to generate an RH-seq reagent for yeast incorporating barcoded transposons (Wetmore et al. 2015). For this purpose, we first generated a pool of plasmids, each encoding a barcoded copy of the piggyBac transposon and its transposase (Figure S1A-C). To use these in RH-seq, we revisited our previously characterized model system: a comparison between DVBPG1373, a thermotolerant Dutch soil strain of *S. cerevisiae,* and Z1, an *S. paradoxus* isolate from the UK (Weiss et al. 2018; AlZaben et al. 2021; Abrams et al. 2021). The F1 hybrid formed from the mating of these strains exhibits a thermotolerance phenotype intermediate between those of the two species parents, and thus is well-suited to mapping of allelic effects on the trait (Weiss et al. 2018). We transformed this F1 hybrid with barcoded plasmids, yielding a pool of hemizygote mutants, which we expanded and then banked (Figure S1D). Next, to catalog the genomic locations of transposon insertions, we used the DNA from a culture of the pool in standard conditions as input into a first round of sequencing library construction, whose primers recognized a common site on the transposon and a common DNA adapter ligated to DNA fragment ends (“Tn-seq”; Figure 1A). Sequencing and data analysis, with quality controls to eliminate barcodes that could not be uniquely associated with a single transposon insertion location (see Methods), yielded a catalog of 548,129 barcoded hemizygotes in the pool whose genomic insertion locations were tabulated.

**Figure 1.**
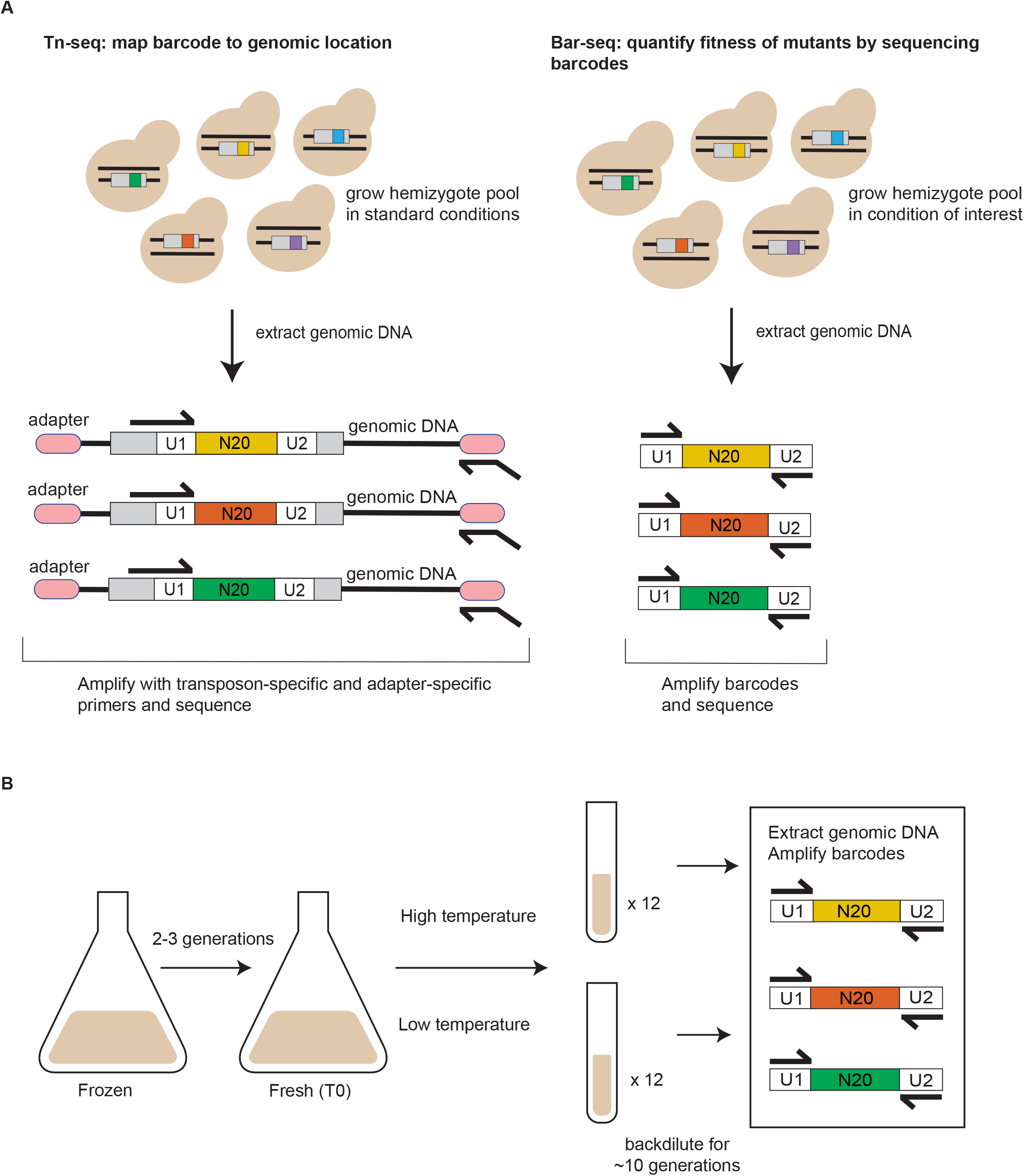
Barcoded RH-seq mapping of yeast thermotolerance loci. (A) Barcoded RH-seq sequencing analysis steps. Left, in a pool of *S. cerevisiae* x *S. paradoxus* hybrid hemizygotes, each harboring a transposon (grey rectangle) marked with a unique 20-mer barcode (multicolored) flanked by universal primer sites (U1 and U2), each barcode is associated with its insertion location by transposon sequencing (Tn-seq). Genomic DNA from the pool is extracted, sheared, and ligated to universal adapters (pink ovals), followed by PCR amplification with a transposon-specific primer (forward black arrow) and an adapter-specific primer (reverse black arrow) and sequencing. Right, for barcode sequencing (Bar-seq) to quantify hemizygote strain abundance after pool growth in a condition of interest, genomic DNA is used as input to PCR with primers to universal primer sites for sequencing. (B) Thermotolerance RH-seq screen design. An aliquot of the hemizygote pool was thawed and cultured in large format, then split into small replicate cultures, each maintained in logarithmic growth phase at the temperature of interest by back-dilution, followed by quantification by Bar-seq.

At this point we could harness the pool for highly replicated screens, each of which could quantify hemizygote abundance in a condition of interest via relatively cheap and straightforward barcode sequencing (“Bar-seq”; Figure 1B).

Thus, with our barcoded hemizygote pool, we implemented an RH-seq screen to search for genes at which *S. cerevisiae* and *S. paradoxus* alleles drove differences in strain abundance at high temperature. For this, we subjected the pool to growth assays with 12 biological replicate cultures at 37°C, alongside controls at 28°C. We used DNA from each culture as input into barcode sequencing (Figure 1B). The resulting data revealed a total of 301,349 cases where a barcode, representing a hemizygote clone with a transposon insertion catalogued by Tn-seq (Figure 1A), was detectable in our growth assays. Transposon insertion positions corresponding to these informative barcodes were evenly split between *S. cerevisiae* and *S. paradoxus* alleles of genes throughout the F1 hybrid genome (Figure S3). We took the normalized count of a given barcode in a sequencing data set as a report of the fitness of the respective hemizygote, *i.e.* its relative abundance after growth in the pool in the respective condition. We then used the complete set of such counts as the input into reciprocal hemizygosity tests to compare, for a given gene, the temperature-dependent abundance of strains harboring a disruption in the *S. cerevisiae* allele, relative to that of strains with the *S. paradoxus* allele disrupted. A pipeline for these tests, including filters for coverage and reproducibility and multiple testing correction (see Methods), revealed 83 genes at a 5% false discovery rate (Figure 2 and Table S7). This contrasted with the much smaller set of eight genes at which species’ alleles drove differences in high-temperature growth, in our original non-barcoded RH-seq approach (Weiss et al. 2018), which had involved only three biological replicates. The 10-fold increase in the number of significant hits in our barcoded RH-seq screen reflects the statistical power afforded by our highly-replicated method to detect even quite small effects.

**Figure 2.**
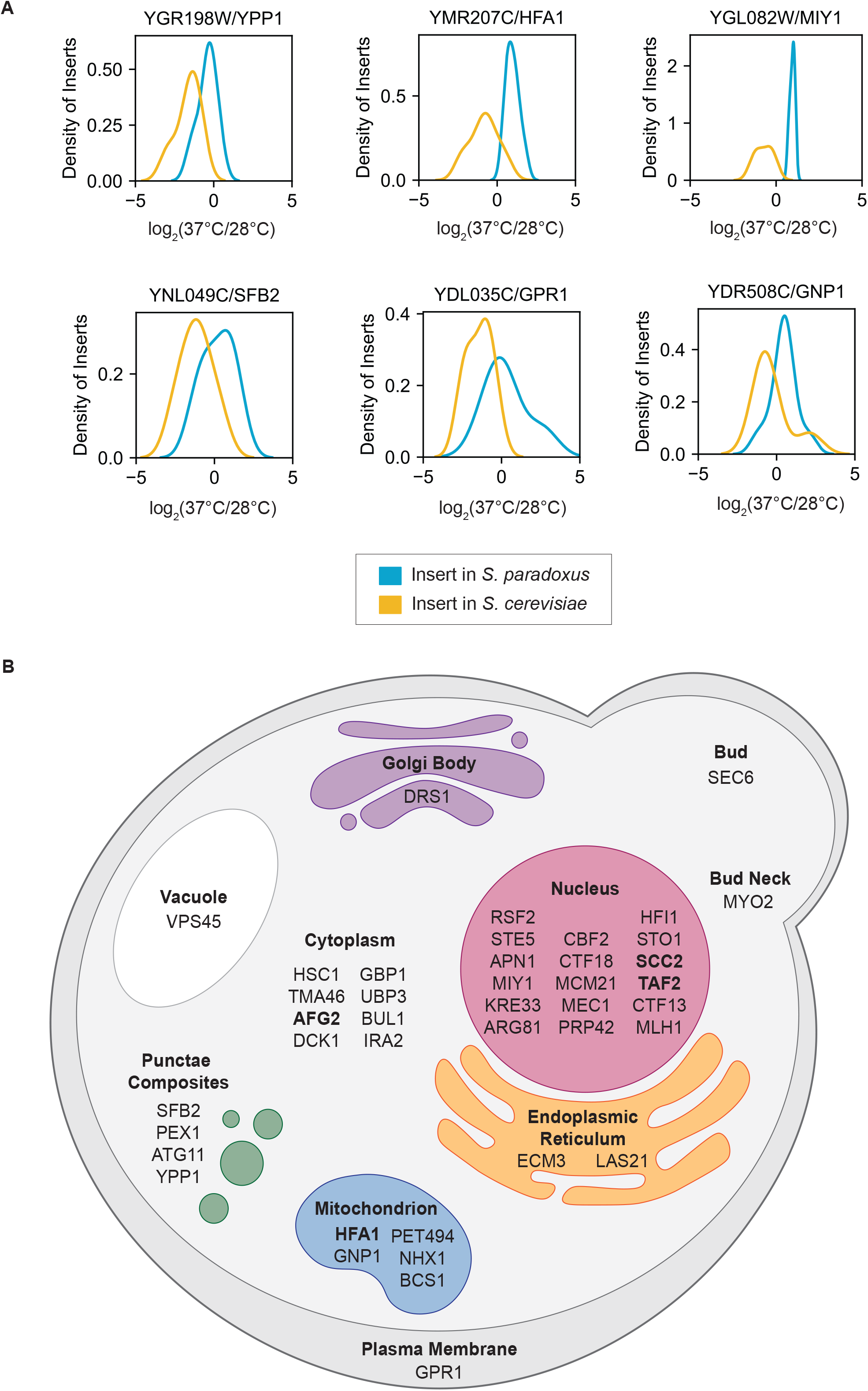
Hits from barcoded RH-seq mapping of yeast thermotolerance. (A) Each panel reports barcoded RH-seq results for a gene at which the *S. cerevisiae* allele is associated with better thermotolerance than the *S. paradoxus* allele, when uncovered in the hybrid background. In a given panel, the *x*-axis reports the log_2_ of abundance, measured by RH-seq after selection at 37°C, of a clone harboring a barcoded transposon insertion in the indicated species’ allele in a given replicate, as a difference from the analogous quantity for that clone after selection at 28°C on average across replicates. The *y*-axis reports the proportion of observations of all clones bearing insertions in the indicated allele that exhibited the abundance ratio on the *x*, as a kernel density estimate. Shown are the top six genes from among all barcoded RH-seq hit loci in terms of allelic effect size; see Table S7 for effect sizes of the complete set of hits. (B) Subcellular localization of RH-seq hit genes, where available from (Pierleoni et al. 2007) and (Huh et al. 2003). Genes at which effects of allelic variation on thermotolerance were reported previously (Weiss et al. 2018; Li et al. 2019) are denoted in bold type.

In our barcoded RH-seq screen hits, as a positive control we first examined the set of genes known to contribute to thermotolerance divergence from our earlier study (*AFG2, APC1, CEP3, DYN1, ESP1, MYO1, SCC2*, and *DYN1*) (Weiss et al. 2018). Several did not meet the experiment-wide statistical thresholds of our barcoded RH-seq pipeline (Figure S4A), suggesting an appreciable false negative rate of the latter overall. However, manual inspection made clear that hemizygosity effects at all gold-standard thermotolerance loci were borne out: in each case, in barcoded RH-seq data, strains with disruptions in the *S. cerevisiae* allele, and a wild-type copy of the *S. paradoxus* allele, had worse thermotolerance than did strains with only the *S. cerevisiae* allele intact (Figure S4A-B), as we had previously reported (Weiss et al. 2018). Furthermore, the list of gene hits from barcoded RH-seq also included *HFA1* (Figure 2B and Tables S7 and S9) which was reported and validated separately as a determinant of thermotolerance differences between yeast species (Li et al. 2019). On the strength of these controls, we considered our deep sampling of thermotolerance loci to serve as a useful proof of concept for the barcoded RH-seq method.

### Functional-genomic analysis of thermotolerance genes

We next aimed to pursue deeper analyses of the novel gene hits from barcoded RH-seq in our yeast thermotolerance application. We considered that a focus on the strongest and most evolutionarily relevant sources of mapping signal would likely yield the most informative results. As such, in light of our interest in explaining the exceptional thermotolerance of purebred *S. cerevisiae*, we earmarked the 44 genes from our larger candidate set at which the *S. cerevisiae* allele boosted the trait most dramatically relative to that of *S. paradoxus* (Figure 2 and Table S9). In what follows, we refer to these genes as our top RH-seq hits, and we analyze them as our highest-confidence predictions for factors that nature would have used in evolving the *S. cerevisiae* phenotype.

We sought to use our mapped loci to explore potential functional mechanisms underlying the thermotolerance trait. We hypothesized that *S. cerevisiae* thermotolerance genes could participate in an interacting network, jointly shoring up particular aspects of cell machinery that were critical for growth at high temperature (AlZaben et al. 2021). Consistent with this notion, the STRING database, which collates experimentally detected protein-protein interactions, genetic interactions, and pathway membership (Szklarczyk et al. 2021), inferred multiple interactions among our top genes from barcoded RH-seq, with salient signal involving cell cycle factors (Figure 3). A more focused analysis revealed an enrichment, among our top barcoded RH-seq hits, for protein-protein interactions with one another as tabulated in BioGRID (Oughtred et al. 2021), to an extent beyond the null expectation (resampling *p* = 0.014). We also implemented qualitative gene set enrichment tests, which revealed that chromosome segregation and mitosis factors, although relatively few in number among our top barcoded RH-seq hit loci, were significantly enriched relative to the genomic null (Table 1). And we developed a complementary, quantitative test to screen Gene Ontology terms for large allelic effect size (the impact on thermotolerance when the *S. cerevisiae* allele of a given gene was disrupted in the hybrid, as a difference from the analogous quantity for the *S. paradoxus* allele; see Methods). Top-scoring in this test was a mitosis gene group, encoding components of the septin ring (GO:0000921; resampling *p* < 0.0001). Together, these results suggest that our top thermotolerance gene hits share commonalities in function, most notably involving cell cycle factors. This dovetails with previous phenotypic and genetic characterization of yeast thermotolerance, including the breakdown of cell division in heat-treated *S. paradoxus* (Weiss et al. 2018), and supports a model in which *S. cerevisiae* acquired thermotolerance in part by resolving the latter cell cycle defect.

**Figure 3.**
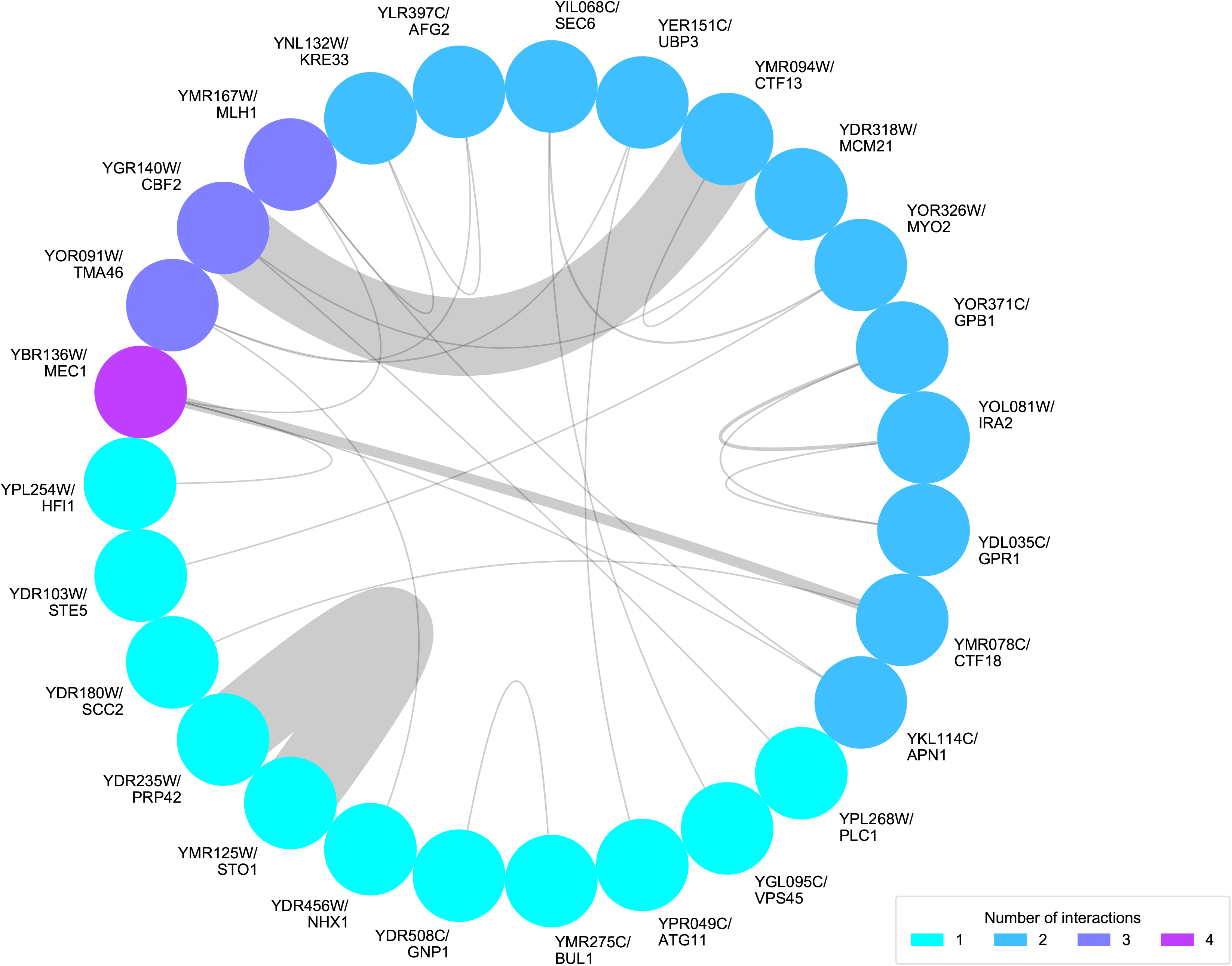
Interactions between thermotolerance loci. Each node represents a top hit gene from barcoded RH-seq mapping of thermotolerance. Each chord represents an inferred interaction, taking into account physical and genetic interactions as well as pathway membership, from the STRING database (Szklarczyk et al. 2021). Chords are weighted by the confidence of the inference of interactions; nodes are colored by the number of inferred interactions with other top hits, such that genes with higher numbers of interactions among the hits are represented by warmer colored nodes.

The genetics of yeast thermotolerance likely also involves mechanisms beside mitosis, given the known role of mitochondrial genes (Baker et al. 2019; Li et al. 2019) and those operating during stationary phase (AlZaben et al. 2021). Indeed, functional-genomic tests revealed enrichment for secretion genes and for regulatory factors in our top RH-seq hits, although no such group constituted a large proportion of the total hit list (Table 1). Annotations in transcription and translation, mitochondrial function, and signaling were also apparent in our top thermotolerance loci (Figure 2B). These trends are consistent with a scenario in which evolution built the trait in *S. cerevisiae* by tweaking an array of housekeeping mechanisms, beside those that involve cell cycle machinery.

### Evolutionary analysis of thermotolerance genes

We anticipated that sequence analyses of the genes we had mapped to thermotolerance by barcoded RH-seq could shed light on the evolutionary history of the trait. To explore this, we used a phylogenetic approach in *Saccharomyces sensu stricto*. We first inferred species-specific branch lengths in the phylogeny of each gene in turn, and focused on the lineage leading to *S. cerevisiae*. The distribution of branch lengths along this lineage among top thermotolerance gene hits was not detectably different from that of the genome as a whole, with the exception of two rapidly evolving thermotolerance genes, *TAF2* and *BUL1*, encoding a transcription initiation factor and ubiquitin ligase adapter respectively (Figure S5). Separately, we quantified protein evolutionary rates in top hits from barcoded RH-seq. A branch-site phylogenetic modeling approach (Yang 2007) detected striking evidence for positive selection along the *S. cerevisiae* lineage in the amino acid permease *GNP1*, the kinetochore DNA binding factor *CBF2*, and the sister chromatid cohesion factor *CTF18* (Figure 4). Interestingly, however, McDonald-Kreitman tests (McDonald and Kreitman 1991) on population-genomic data did not detect an overall excess of amino acid variation relative to synonymous changes, at these three genes or any other barcoded RH-seq hit locus (Table S10). Thus, even at genes harboring individual codons with likely signatures of selection, we could not detect evidence for a scenario where *S. cerevisiae* stacked up a large number of unique amino acid changes, in the evolution of thermotolerance. Together, however, our analyses do highlight thermotolerance genes with marked signal for derived alleles in *S. cerevisiae* at single codons or in the overall DNA sequence—cases where species divergence is likely to be of phenotypic and evolutionary importance.

**Figure 4.**
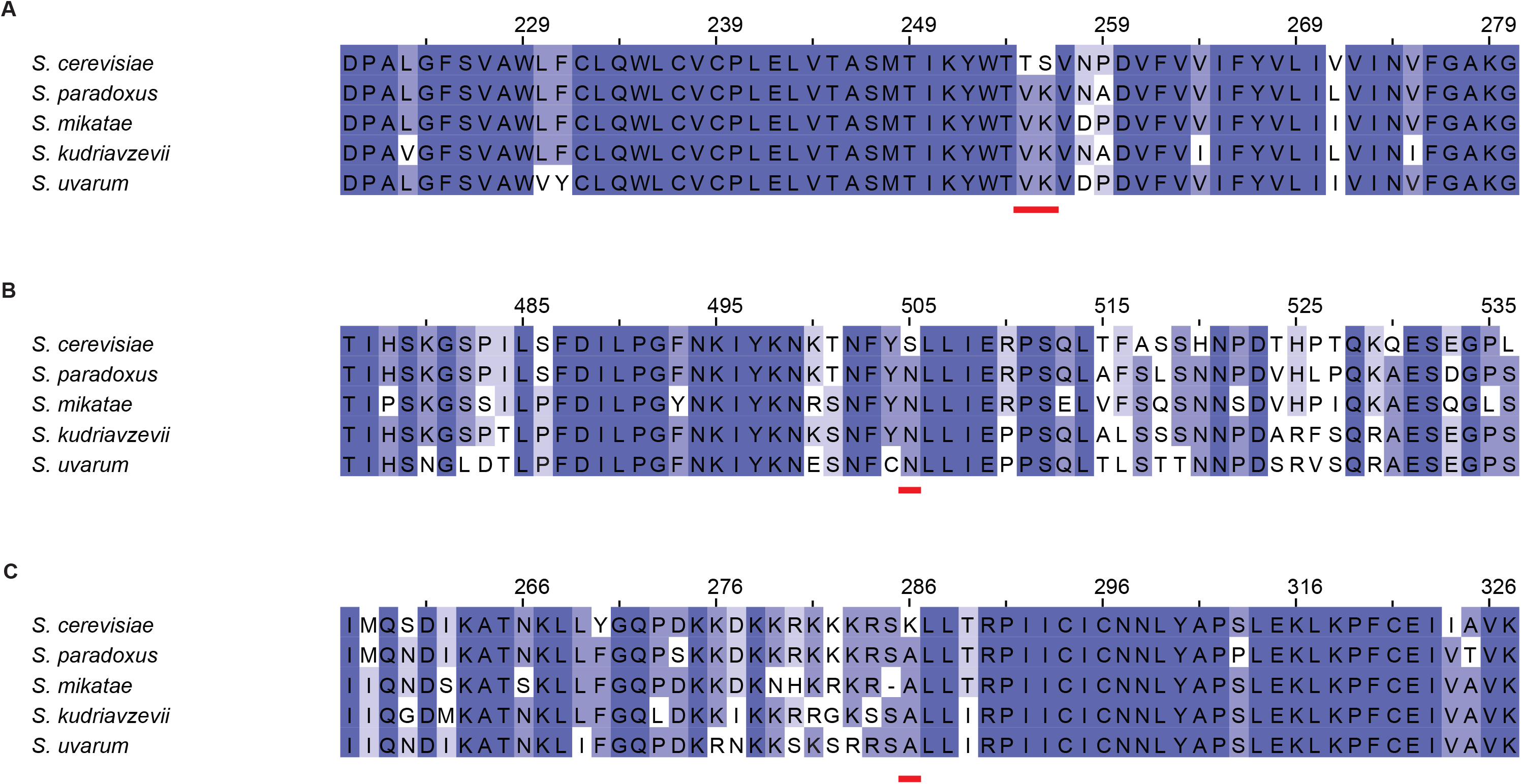
Codons under positive selection in thermotolerance loci. Each panel shows the amino acid sequence context of the codon(s) (red bar) inferred to be under positive selection along the *S. cerevisiae* lineage, in a hit gene from RH-seq thermotolerance mapping. Alignments are colored by percent identity, with darker purples indicating a higher percent identity. (A) *YDR508C/GNP1*. (B) *YGR140W/CBF2*. (C) *YMR078C/CTF18*.

## DISCUSSION

### RH-seq power and the interpretation of mapped loci

In this work, we established the barcoded RH-seq method for genetic dissection of trait variation between diverged lineages. RH-seq falls into a family of recently-developed methods that can dissect natural trait variation across species barriers (Weiss and Brem 2019). A chief distinction of RH-seq is its low cost and low overhead, and the barcoding feature we add here cuts down labor and cost even further, enabling high replication.

Our application to yeast thermotolerance serves as an informative model for the performance of barcoded RH-seq on highly genetically complex traits. We pinpointed dozens of candidate genes at which species-level variation contributes to growth at high temperature. And yet we also observed evidence for a sizeable false negative rate among our barcoded RH-seq results, since some validated thermotolerance loci from our earlier screen did not appear among the hits here. Likewise, a separate barcoded RH-seq mapping of yeast species’ differences in growth under milder heat stress revealed little signal above the noise (Table S6 and Table S8), likely reflecting very weak genetic effects under this condition. We thus expect that, as would be true for a classical linkage or association scan, the statistical power of a barcoded RH-seq experiment is a function of signal to-noise, genetic complexity, and genetic effect size; and that many thermotolerance loci remain to be identified even in our very deep set of screen results from high-temperature growth.

By virtue of our focus on pro-thermotolerance alleles in *S. cerevisiae*, our work has left open the functional and evolutionary genomics of loci at which the allele from *S. cerevisiae* instead conferred worse thermotolerance than that of *S. paradoxus*, when each in turn was uncovered in the hybrid. Our barcoded RH-seq identified a number of such genes at high statistical significance. These loci may well reflect the accumulation of advantageous alleles in *S. paradoxus*, or deleterious alleles in *S. cerevisiae*, by drift, even as the latter was under selection to improve the trait in evolutionary history. Analogously, in linkage mapping results, the effect of an allele in recombinant progeny from a cross often does not conform to that expected from the respective parent’s phenotype (Burke and Arnold 2001; Brem and Kruglyak 2005). It is also possible that some such allelic effects are the product of epistatic interactions between a locus of interest and the hybrid background, and would be phenotypically buffered (and thus evolutionarily irrelevant) in the purebred species. Molecular validation will be necessary to confirm the phenotypic impact of variation at our mapped loci, and its potential dependence on genetic background.

That said, we consider genes with pro-thermotolerance *S. cerevisiae* alleles according to barcoded RH-seq to be strong candidates for *bona fide* determinants of the trait from the wild in this species. Indeed, earlier work has shown that for such genes mapped by RH-seq in the hybrid, the advantage of *S. cerevisiae* alleles is borne out in tests in purebred backgrounds (Weiss et al. 2018). Accordingly, we have shown here that as a cohort, barcoded RH-seq hits with advantageous *S. cerevisiae* alleles exhibit functional and sequence-based attributes consistent with a role in thermotolerance evolution in the wild.

### Cellular and molecular mechanisms of thermotolerance

Our top RH-seq hits revealed strong evidence for chromosome segregation and other mitosis functions as a linchpin of *S. cerevisiae* thermotolerance. As a complement to earlier characterization of six such genes (*APC1, ESP1, DYN1, MYO1, CEP3,* and *SCC2*) (Weiss et al. 2018; Abrams et al. 2021), we now report seven new thermotolerance determinants that function in cell division (*MEC1, MLH1, CTF13, CTF18, MCM21, CBF2,* and *MYO2*). The emerging picture is one in which the ancestor of modern-day *S. cerevisiae*, faced with dysfunction of a slew of mitotic factors at high temperature, acquired variants across the genome to shore up their activity under these conditions. Under one model of *S. cerevisiae* evolution, the particular niche to which this species specialized was one of avid fermentation, producing (and resisting) heat and ethanol at levels that eliminated its microbial competitors (Goddard 2008; Salvadó et al. 2011). In such a scenario, the maximum benefit could well accrue to the organism if it were able to undergo rapid cell division under the challenging conditions of its own making. Consistent with this notion, another budding yeast, *Hanseniaspora,* which often dominates in early fermentation prior to takeover by *S. cerevisiae* (Fleet 2003), underwent evolutionary loss of much of the cell-cycle checkpoint machinery, consistent with a strategy of accelerated growth at any cost to outcompete other species at the respective stage (Steenwyk et al. 2019).

However, since our current hit list includes many genes from other housekeeping pathways, from transcription/translation to transport and lipid metabolism, mitosis does not appear to be the whole mechanistic story for the thermotolerance trait in *S. cerevisiae*. Indeed, other housekeeping factors also showed up in our previous screen (Weiss et al. 2018) and in an elegant complementary study of mitochondrial determinants of thermotolerance divergence between yeast species (Baker et al. 2019; Li et al. 2019). The panoply of functions detected among our mapped loci conforms well to current models of the mechanisms of thermotolerance, which invoke many essential genes and housekeeping processes (Leuenberger et al. 2017).

The latter idea emerged largely from a proteomic study which showed that thermotolerant organisms had higher thermostability of essential proteins of many functions, across the tree of life (Leuenberger et al. 2017). Were sequence changes that led to improved protein stability a linchpin of thermotolerance evolution in *S. cerevisiae*? Our data are consistent with a mechanistic role for properties of the protein sequences of many thermotolerance genes, in that variation in coding regions has come to the fore in our sequence tests here and those of an earlier small-scale analysis (Abrams et al. 2021). And interestingly, an experimental case study of one of our mapped thermotolerance loci revealed no impact on the trait from variation in the promoter, only from that in the coding region (Abrams et al. 2021). We cannot rule out noncoding determinants in some cases, especially given that a few hundred genes exhibit temperature-dependent *cis*-regulatory programs unique to *S. cerevisiae* (Tirosh et al. 2009; Li and Fay 2017). But if coding regions do hold the exclusive key to the mechanism of *S. cerevisiae* thermotolerance, they could well involve variants that improve protein function and regulation alongside folding/structure at high temperature. Overall, then, we envision that nature could have used a variety of molecular mechanisms in building the trait, given the apparent complexity of the problem. Biochemical studies will be necessary to nail down exactly how *S. cerevisiae* alleles advance thermotolerance.

In summary, our data reveal a newly detailed picture of the highly polygenic architecture for a natural trait divergence between species. It is tempting to speculate that evolution may draw on a vast number of variants across the genome to refine a trait over millions of generations, making effects stronger, weaker, or less pleiotropic, adding regulatory control, and so on (Orr 1998). If so, these architectures may ultimately conform to the omnigenic model (Boyle et al. 2017)–which was originally applied to human disease genetics, but may also prove to be an apt description of ancient adaptations.

## ACKNOWLEDGEMENTS

This work was supported by NSF GRFP DGE 1752814 to M.A. and NIH R01 GM120430 to R.B.B. The authors thank Adam Arkin for his generosity with computational resources; Morgan Price, Lori Huberman, and members of the John Dueber lab for discussions about barcoded transposon mutagenesis; and sequencing; and Abel Duarte for cloning advice.

## SUPPLEMENTARY FIGURE CAPTIONS

**Supplementary Figure 1. Making barcoded hemizygotes in a yeast hybrid background for RH-seq.** (A) A pool of random N20 barcodes (colors), each flanked by universal priming sites (U1 and U2), was used as input into a PCR with primers containing recognition sites for the BbsI type IIS restriction enzyme. (B) In a plasmid harboring an un-barcoded piggyBac transposon (gray rectangle) (the kanamycin resistance cassette, kan^R^, flanked by left and right transposon arms) and transposase (teal rectangle) (Weiss et al. 2018), a 42 nucleotide stuffer sequence, consisting of two BbsI restriction enzyme sites flanking a NotI restriction enzyme site and custom overhang sequences (Lee et al. 2015), replaced 42 nucleotides in the right arm of the transposon. (C) BbsI digestion of the barcodes and stuffer-containing plasmid, followed by stuffer loss and ligation of a barcode into each plasmid, yielded a pool of barcoded plasmids. (D) Transformation of the barcoded transposase plasmid into *S. cerevisiae* x *S. paradoxus* hybrids, followed by transposition and plasmid loss, yielded a pool of marked transposon hemizygote insertion genotypes in the hybrid background.

**Supplementary Figure 2. Modifying yeast piggyBac to test barcode insertion positions and transposase optimization, and effects of barcode insertion positions and transposase sequence on yeast piggyBac transposition efficiency.** (A) Left, a plasmid from (Weiss et al. 2018) containing the unbarcoded piggyBac transposon (gray) and transposase (teal) was modified to eliminate three BbsI restriction enzyme sites, and used as a backbone for further modifications. Right, test plasmids were mutated at transposase sites designed to optimize codons and increase activity (Yusa et al. 2011). Bottom, test plasmids were modified to incorporate into the transposon a single 20 nucleotide barcode flanked by universal priming regions and custom two-nucleotide overhang sequences (blue squares), either by insertion between the 3’-most end of the left arm and 5’ end of the TEF promoter of the kanamycin cassette (bottom left) or replacing endogenous nucleotides inside the right arm of the transposon (bottom right). Pink rectangles indicate transposase binding sites from (Morellet et al. 2018). (B) Each pair of boxes reports transposition test results from a plasmid schematized in (A) in the *S. cerevisiae* x *S. paradoxus* F1 hybrid, with transformation at the indicated temperature. For a given box, the thick black line reports the median; the box extent report quartiles; whiskers report outliers.

**Supplementary Figure 3. Gene coverage and read depth in thermotolerance Bar-seq.** (A) The *x*-axis reports the number of inferred hemizygote clones in a given gene (corresponding to transposon insertion mutants) whose abundance was detectable in Bar-seq (see Figure 1B), and each bar height reports the number of genes with the number of detectable hemizygotes on the *x*, for the indicated species’ allele in the diploid hybrid. The dotted red line indicates the cutoff used in our quality control pipeline for tests of allelic impact on thermotolerance, whereby only genes with greater than three inserts for an allele in the Bar-seq counts were considered. (B) The *x*-axis reports the total number of Bar-seq reads, for a given inferred hemizygote clone in the indicated species’ allele, in competition cultures grown at 28°C; each bar height reports the number of inferred hemizygote clones with the Bar-seq abundance on the *x*. (C) Data are as in (B) except that competitions at 37°C were analyzed.

**Supplementary Figure 4. Impact on high-temperature growth of allelic variation, in barcoded RH-seq, at genes from a previous thermotolerance screen.** (A) Each row reports the allelic effect, the thermotolerance conferred by disruption of the *S. cerevisiae* allele, relative to the analogous quantity for the *S. paradoxus* allele, as measured in barcoded RH-seq, of a gene at which allelic variation was previously reported to impact thermotolerance (Weiss et al. 2018). Genes marked with asterisks were significant at *p* < 0.05, after quality control for noise and number of inserts and multiple-hypothesis correction. (B) The *x*-axis reports allelic effect for a given gene as in (A); the *y*-axis reports the proportion of genes with the allelic effect on the *x*, with the blue trace showing the distribution across all genes with barcoded RH-seq data, as a kernel density estimate. Red dotted vertical lines represent genes from (A).

**Supplementary Figure 5**. **Accelerated evolution of thermotolerance loci.** Shown are results of analyses of branch length of top hit genes from barcoded RH-seq mapping of thermotolerance, as inferred from gene trees and normalized for gene length. Each vertical bar reports inferred branch length, along the *S. cerevisiae* lineage, of the indicated RH-seq hit gene. Horizontal lines report median branch lengths across the indicated gene sets. A resampling test for long branches on the *S. cerevisiae* lineage among top RH-seq hits revealed significant evidence for enrichment (*p* = 0.0465) but not when *TAF2* and *BUL1* were eliminated (*p* = 0.1574), attesting to the particularly strong inference of accelerated evolution in the latter two genes.

## SUPPLEMENTARY TABLES

**Supplementary Table 1.**
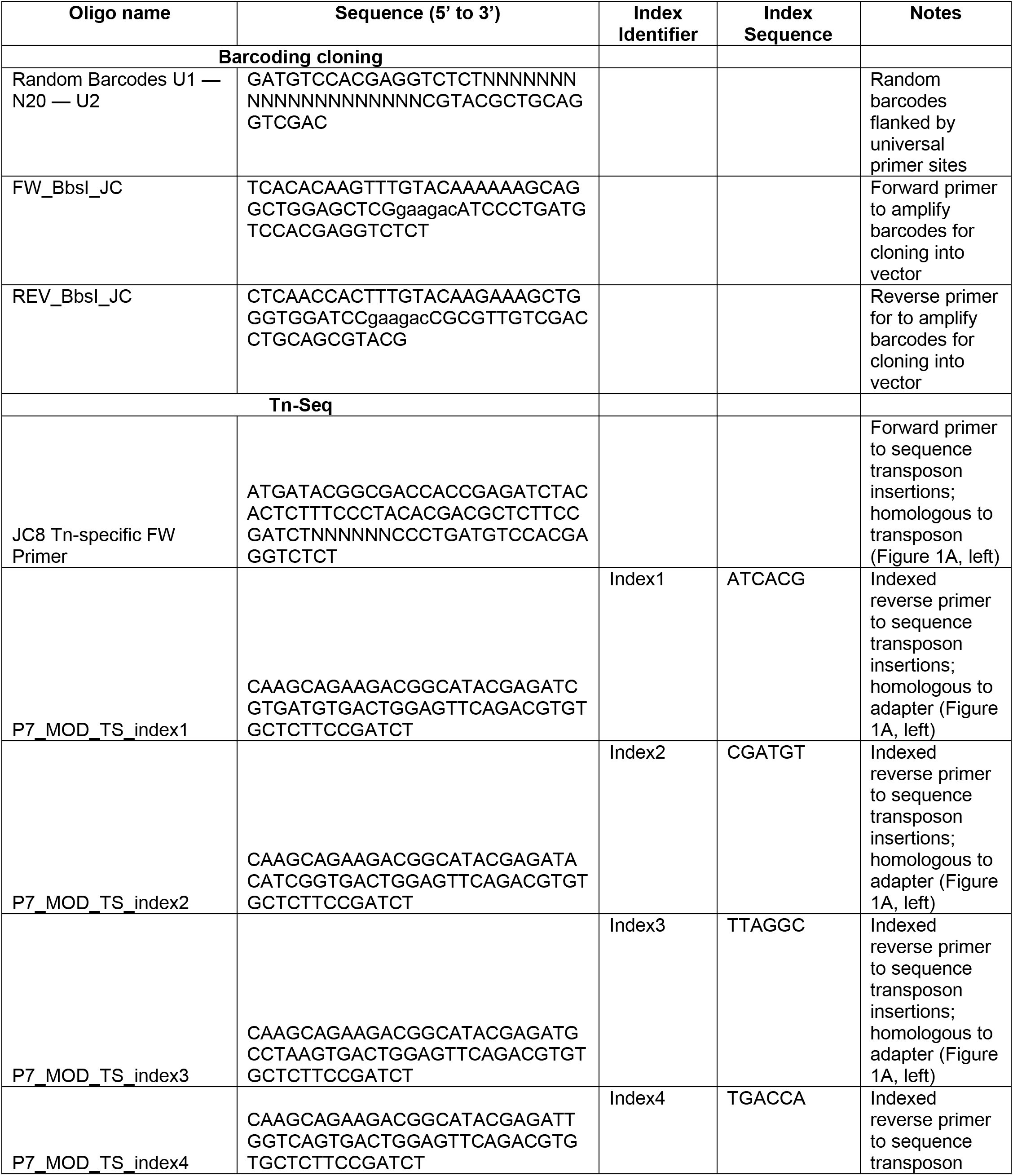
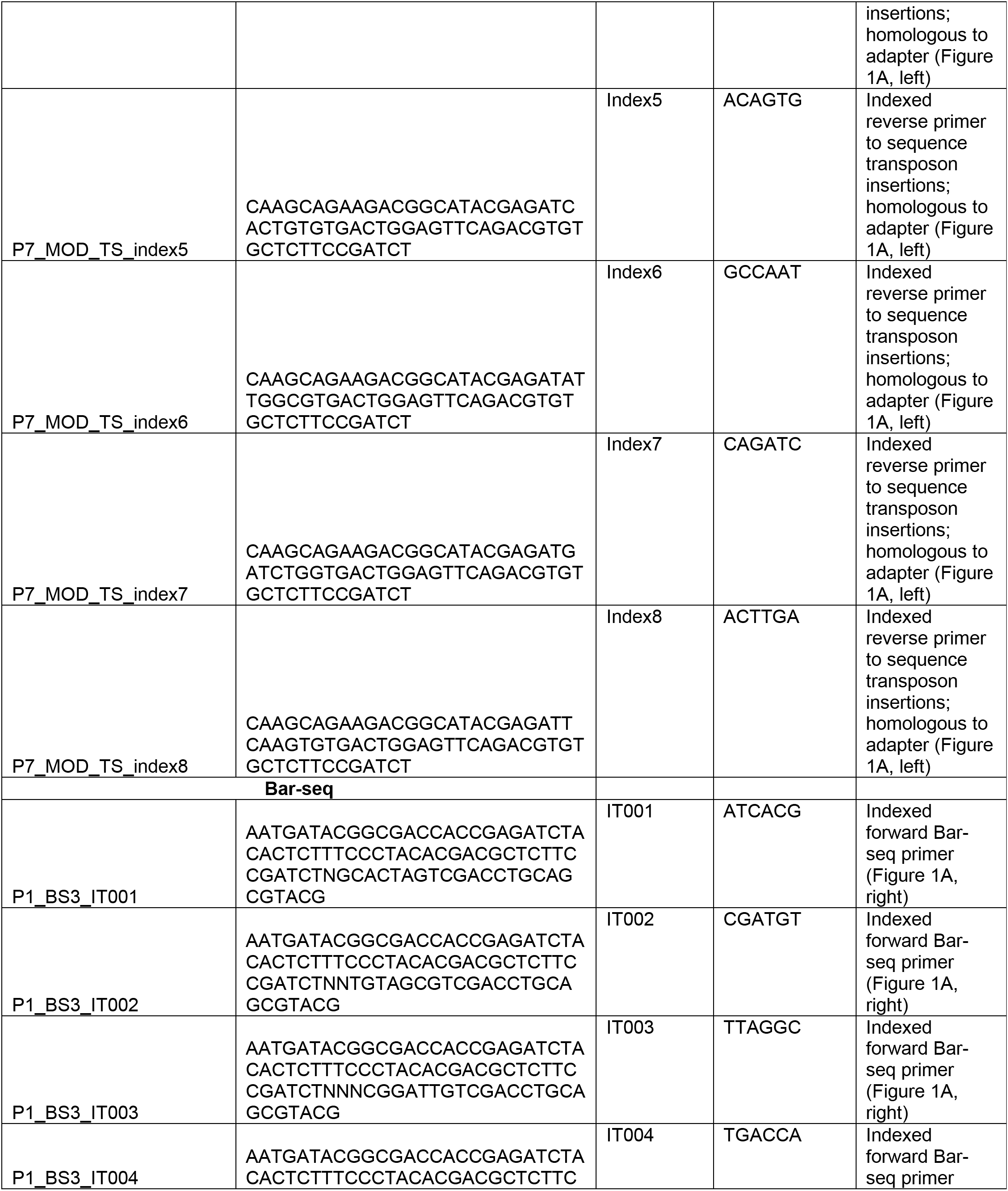
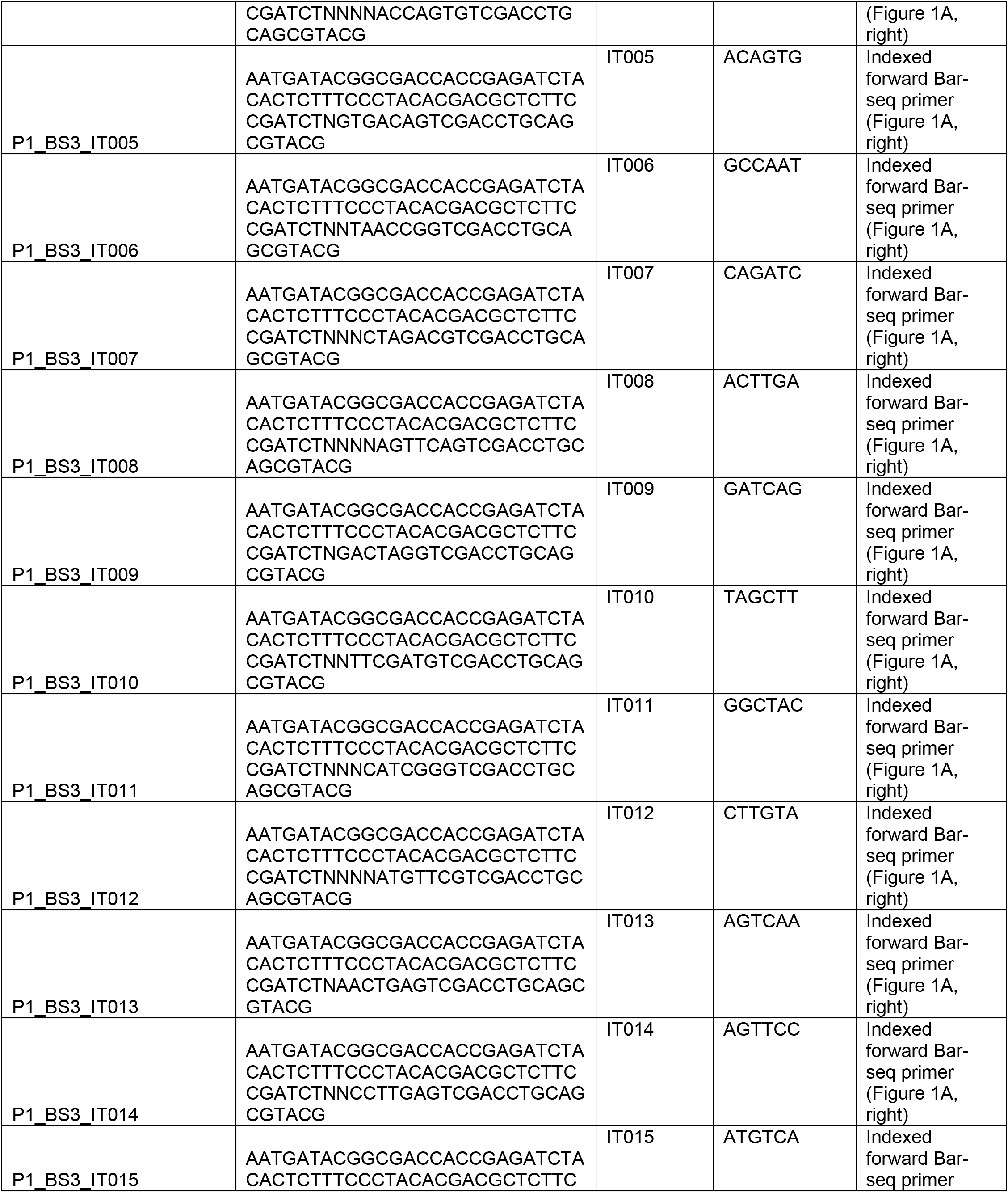
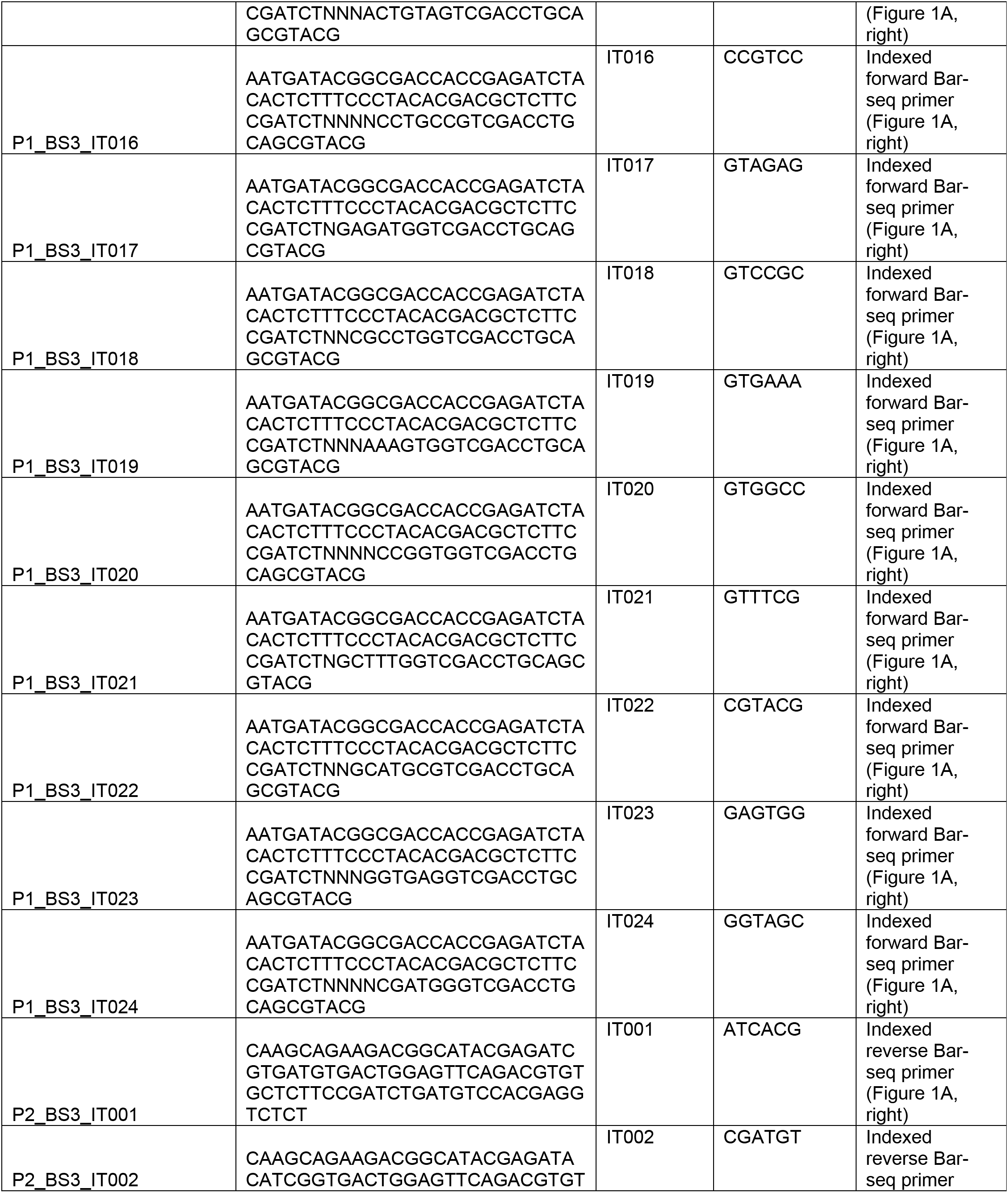
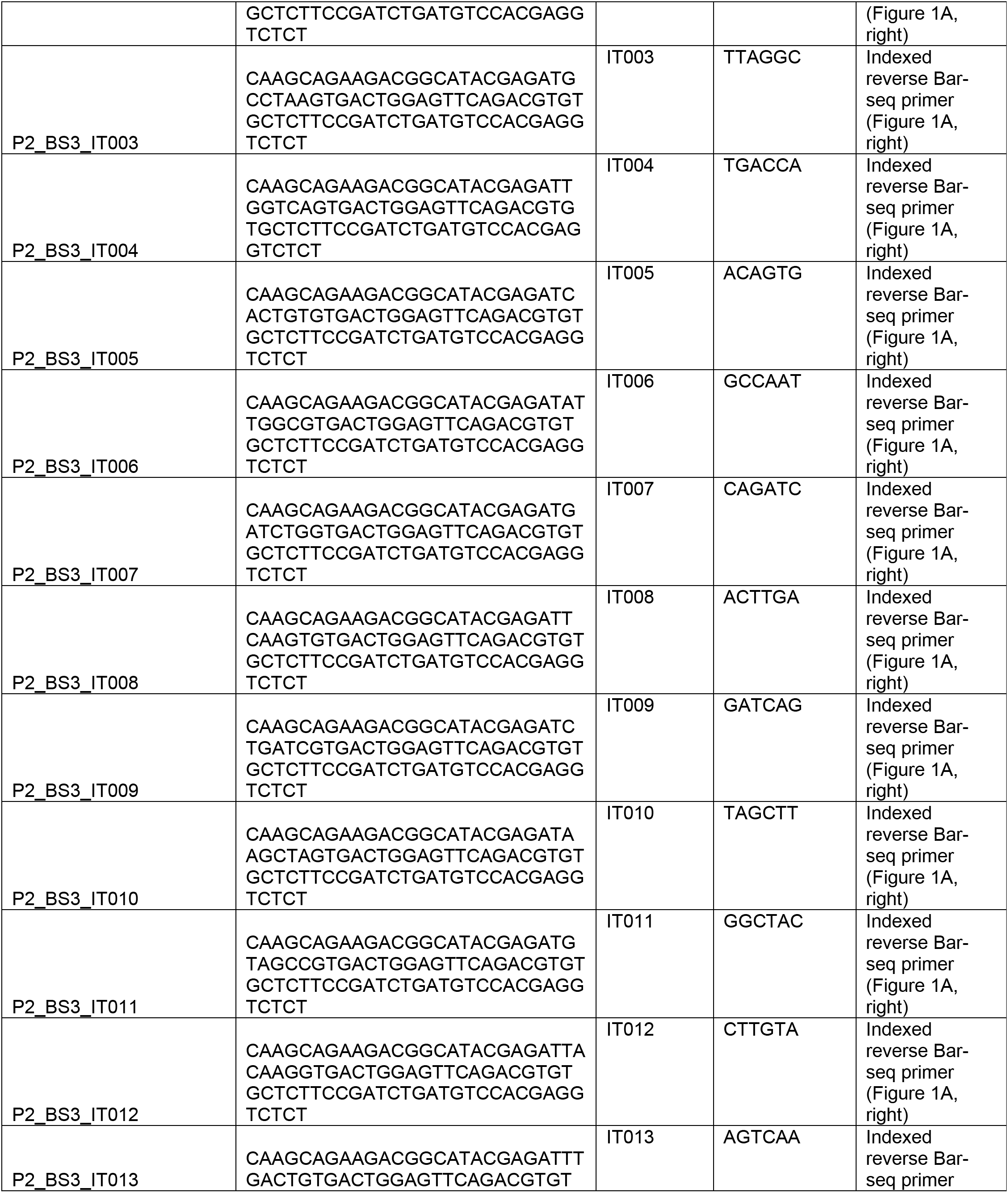
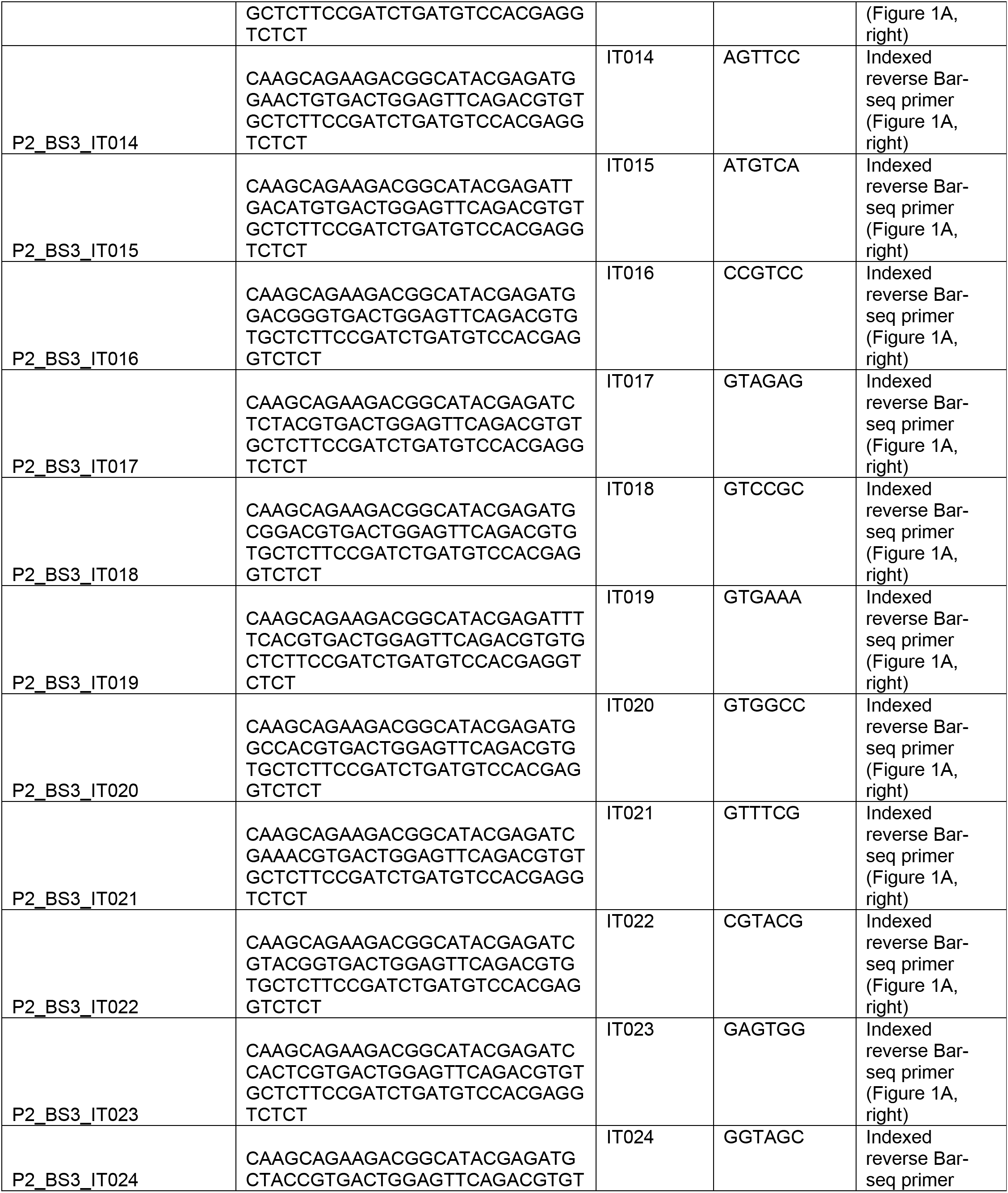
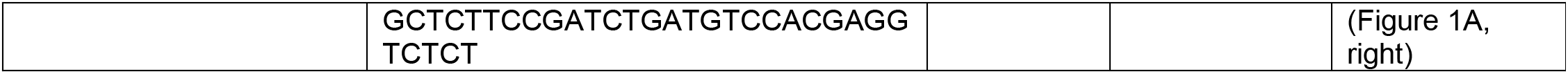
Plasmids used in this study.

**Supplementary Table 2.** Oligonucleotides used in this study.

| Experiment | Library | Reads |
| --- | --- | --- |
| *E. coli* vector pool Bar-seq | RBJC009_IT013 | 93,277,156 |
|  | RBJC009_IT014 | 73,914,499 |
|  | RBJC009_IT015 | 79,273,122 |
|  | RBJC009_IT016 | 81,404,878 |
|  | RBJC45_IT013 | 100,794,463 |
|  | RBJC45_IT014 | 92,410,111 |
|  | RBJC45_IT015 | 91,931,624 |
|  | RBJC45_IT016 | 96,631,037 |
|  | RBJC009_IT013 | 93,277,156 |
|  | RBJC009_IT014 | 73,914,499 |
|  | RBJC009_IT015 | 79,273,122 |
|  | RBJC45_IT016 | 96,631,037 |
| 37°C vs. 28°C barcoded RH-seq in yeast | RBMA038A_IT001 | 16,042,537 |
|  | RBMA038A_IT002 | 19,451,725 |
|  | RBMA038A_IT003 | 16,313,279 |
|  | RBMA038A_IT004 | 19,512,043 |
|  | RBMA038A_IT005 | 13,625,004 |
|  | RBMA038A_IT006 | 22,257,379 |
|  | RBMA038A_IT007 | 22,385,133 |
|  | RBMA038A_IT008 | 16,359,601 |
|  | RBMA038A_IT009 | 20,118,293 |
|  | RBMA038A_IT010 | 19,121,369 |
|  | RBMA038A_IT011 | 18,887,044 |
|  | RBMA038A_IT012 | 15,895,905 |
|  | RBMA038A_IT013 | 18,922,229 |
|  | RBMA038A_IT014 | 26,602 |
|  | RBMA038A_IT015 | 16,571,744 |
|  | RBMA038A_IT016 | 18,783,445 |
|  | RBMA038A_IT017 | 17,671,257 |
|  | RBMA038A_IT018 | 18,040,605 |
|  | RBMA038A_IT019 | 16,929,985 |
|  | RBMA038A_IT020 | 15,319,721 |
|  | RBMA038A_IT021 | 16,897,401 |
|  | RBMA038A_IT022 | 16,372,841 |
|  | RBMA038A_IT023 | 16,679,583 |
|  | RBMA038A_IT024 | 13,875,240 |
| 36°C vs. 28°C barcoded RH-seq in yeast | RBMA039A_IT001 | 28,004,341 |
|  | RBMA039A_IT002 | 19,283,097 |
|  | RBMA039A_IT003 | 12,427,040 |
|  | RBMA039A_IT004 | 13,934,236 |
|  | RBMA039A_IT005 | 12,068,707 |
|  | RBMA039A_IT006 | 60,072,532 |
|  | RBMA039A_IT008 | 12,500,340 |
|  | RBMA039A_IT009 | 25,029,516 |
|  | RBMA039A_IT010 | 14,765,007 |
|  | RBMA039A_IT011 | 7,059,133 |
|  | RBMA039A_IT012 | 19,837,690 |
|  | RBMA039A_IT013 | 14,393,602 |
|  | RBMA039A_IT014 | 19,572,649 |
|  | RBMA039A_IT015 | 8,814,383 |
|  | RBMA039A_IT016 | 14,587,181 |
|  | RBMA039A_IT017 | 8,619,661 |
|  | RBMA039A_IT018 | 13,216,843 |
|  | RBMA039A_IT019 | 12,499,486 |
|  | RBMA039A_IT020 | 11,963,866 |
|  | RBMA039A_IT021 | 6,626,989 |
|  | RBMA039A_IT022 | 11,649,542 |
|  | RBMA039A_IT023 | 5,813,357 |
|  | RBMA039A_IT024 | 9,349,172 |

**Supplementary Table 3.** Bar-seq sequencing data sets. Each row reports numbers of reads sequenced for the indicated Bar-seq experiment. The first set of rows reports results from a check of barcoded piggyBac transposon plasmids as in Figure S1C; the remaining rows report results from quantification of yeast hemizygote insertion genotypes after competition in the indicated condition, as in Figure 1B of the main text. Experiment identifiers are from BioProject PRJNA735401.

| Pool | Library | Reads | Platform | Facility |
| --- | --- | --- | --- | --- |
| 67 | RBJC37 | 38,713,102 | Novaseq SP<br>PE150 | UC Berkeley |
| 69 | RBJC38 | 38,875,221 |  |  |
| 69 | RBJC39 | 43,194,450 |  |  |
| 69 | RBJC40 | 39,778,862 |  |  |
| 69 | RBJC41 | 38,836,065 |  |  |
| 67 | RBJC42 | 39,265,466 |  |  |
| 67 | RBJC43 | 47,124,575 |  |  |
| 67 | RBJC44 | 39,762,187 |  |  |
| 67 | RBJC48 | 91,531,071 | HiSeq4000 PE150 | Novogene |
| 67 | RBJC48 reseq | 86,892,060 |  |  |
| 70 | RBCJ51 | 86,254,426 |  |  |
| 70 | RBCJ51 reseq | 86,130,880 |  |  |
| 70 | RBJC52 | 52,108,306 |  |  |
| 70 | RBJC52 reseq | 53,363,169 |  |  |
| 71 | RBJC54 | 88,154,532 |  |  |
| 71 | RBJC54 reseq | 86,878,835 |  |  |
| 71 | RBJC55 | 90,265,981 |  |  |
| 71 | RBJC55 reseq | 82,130,170 |  |  |
| 69 | RBJC57 | 84,296,399 |  |  |
| 69 | RBJC57 reseq | 85,606,080 |  |  |

**Supplementary Table 4. Tn-seq sequencing data sets.** Each row reports numbers of reads from the indicated sequencing of insertion positions of barcoded transposons in the *S. cerevisiae* x *S. paradoxus* hybrid, as in Figure 1A, left, of the main text. Experiment identifiers are from PRJNA735401; “reseq” indicates the reads from a technical replicate performed to gather additional reads for the indicated library.

**Supplementary Table 5. Abundance of inferred hemizygote insertion genotypes from thermotolerance RH-seq.** Each row reports the results of sequencing from one inferred transposon insertion in the *S. cerevisiae* x *S. paradoxus* diploid hybrid after selection of the barcoded transposon pool after competitions comparing growth at 37°C and 28°C, reflecting the abundance in the pool of the respective hemizygote clone harboring the insertion. Chromosome, strand, location, and gene report the fine-scale position of the inferred insertion. Allele, the species parent’s homolog in which the transposon insertion lay. Abundance, read counts of the transposon insertion sequenced after selection of the barcoded transposon pool at the indicated temperature, normalized for library size and averaged across the biological replicate cultures. Transposon insertions not detected in any replicate of the indicated selection were assigned an abundance of 1 prior to normalization by library size. CV, coefficient of variation over biological replicates of normalized read counts after selection at the indicated temperature. Barcode, the unique barcode identifier of the transposon insertion.

**Supplementary Table 6. Abundance of hemizygote insertion genotypes from RH-seq at 36°C.** Data are as in Table S5, except that RH-seq was done using 36°C as the high-temperature condition.

**Supplementary Table 7. Effects of allelic variation in thermotolerance RH-seq.** Each row reports the results of reciprocal hemizygote tests on thermotolerance at the indicated gene in the *S. cerevisiae* x *S. paradoxus* diploid hybrid at 37°C. Columns B-G report analyses of abundance upon the aggregation at the gene level of inferred hemizygote genotypes (Table S5) from all biological replicate experiments, filtered for quality control (see Methods). Columns B-D report results of a two-tailed Mann-Whitney statistical test for a difference in the abundance after growth at 37°C, relative to the abundance after growth at 28°C, of hemizygotes harboring transposon insertions in the two species parents’ homologs. The Benjamini-Hochberg method was used to correct for multiple testing. Column E reports the log_2_(abundance at 37°C/abundance at 28°C) of the average insert in the *S. cerevisiae* allele. Column F reports the analogous quantity among inserts in the *S. paradoxus* allele of the gene. Column G reports the allele-specific effect size, calculated as the difference between the measures of Columns E and F.

**Supplementary Table 8.**
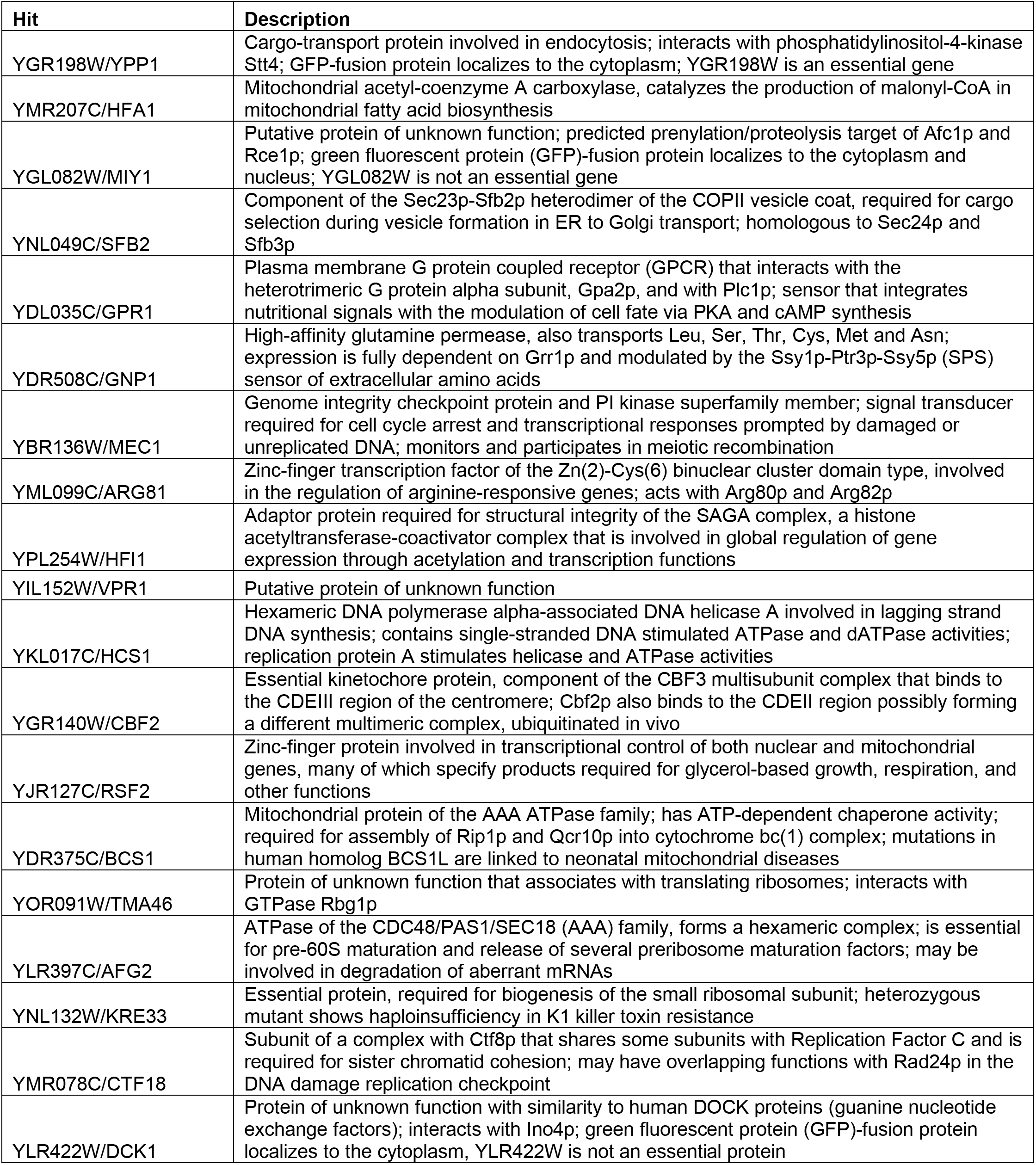
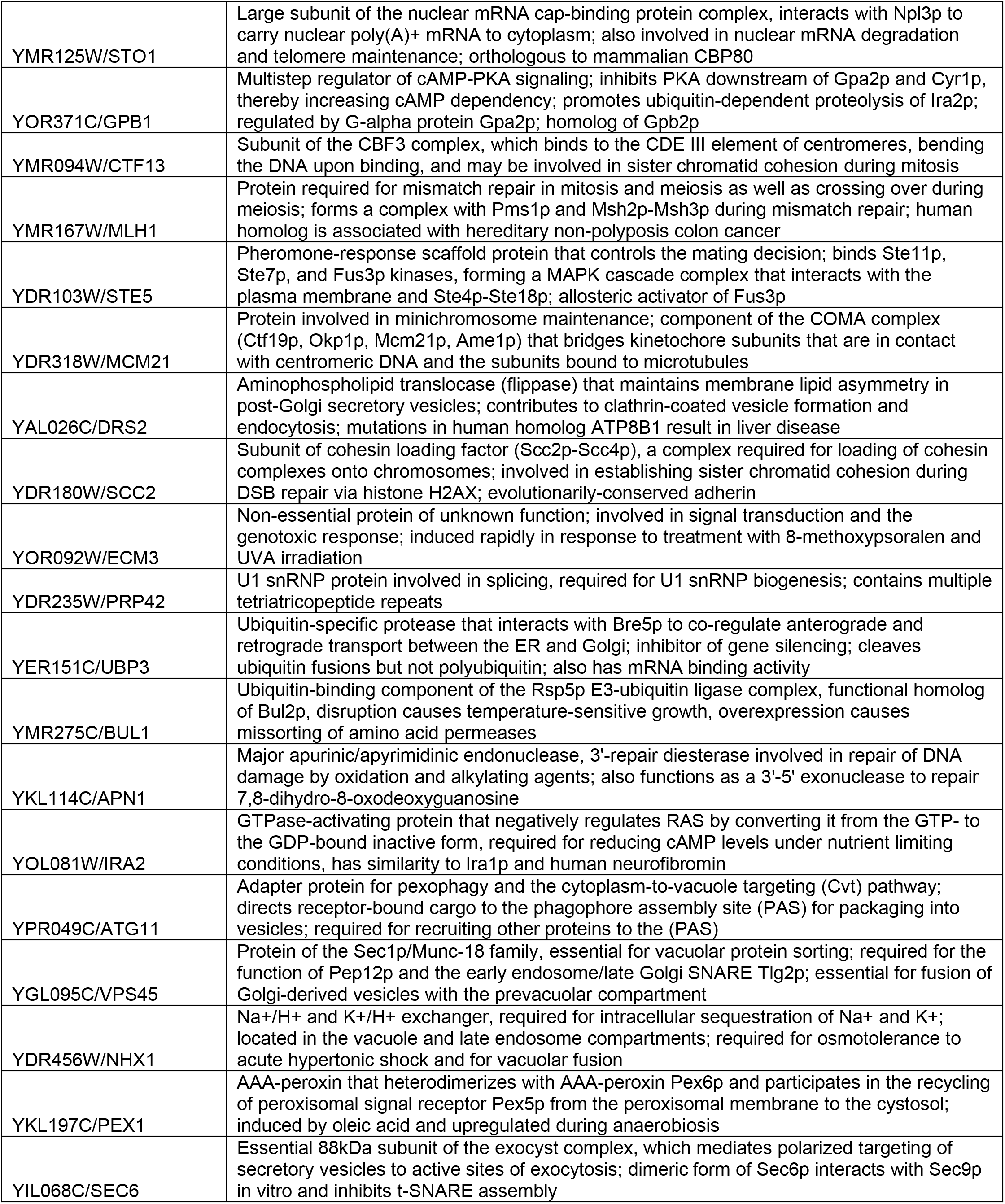
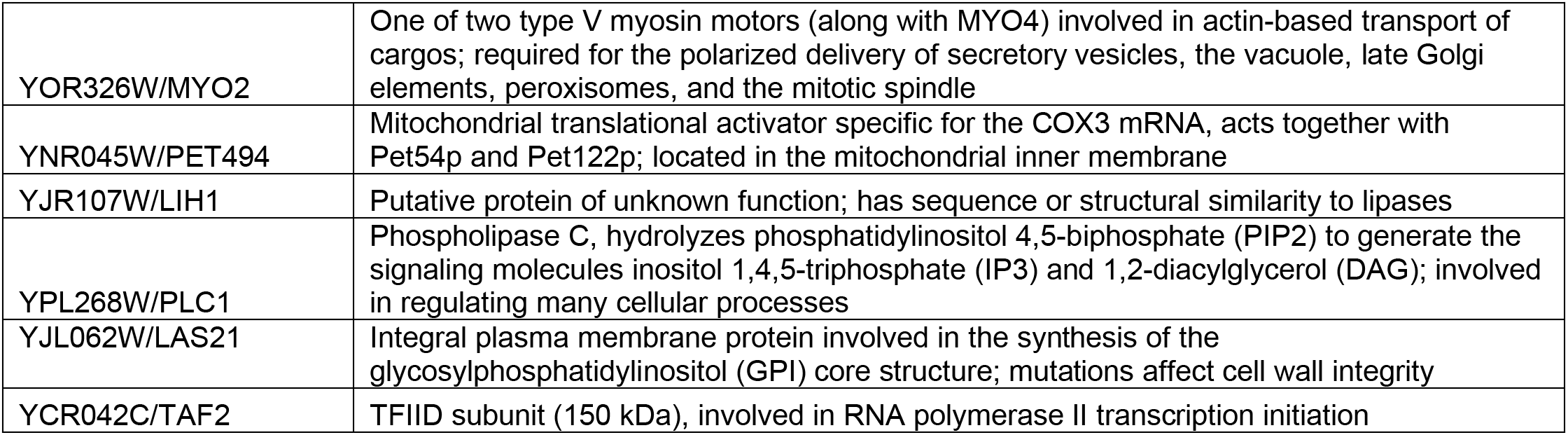
Effects of allelic variation in RH-seq at 36°C. Data are as in Table S7, except that the high temperature growth condition was 36°C.

**Supplementary Table 9.** Annotations of top hit loci from barcoded RH-seq of thermotolerance. Shown are hits from thermotolerance mapping by barcoded RH-seq (Table S7) that met quality control thresholds and at which disruption of the *S. cerevisiae* allele compromised thermotolerance to a greater extent than did disruption of the *S. paradoxus* allele in the interspecific hybrid.

| gene | D<sub>s</sub> | D<sub>n</sub> | P<sub>s</sub> | P<sub>n</sub> | NI | p | adjusted p |
| --- | --- | --- | --- | --- | --- | --- | --- |
| YBR136W/MEC1 | 293 | 38 | 770 | 363 | 3.63 | 5.71 x 10<sup>-15</sup> | 1.69 x 10<sup>-12</sup> |
| YLR422W/DCK1 | 122 | 26 | 335 | 280 | 3.92 | 1.13 x 10<sup>-10</sup> | 8.70 x 10<sup>-9</sup> |
| YOR326W/MYO2 | 35 | 0 | 1486 | 1273 | inf | 5.15 x 10<sup>-10</sup> | 2.77 x 10<sup>-3</sup> |
| YAL026C/DRS2 | 32 | 1 | 1399 | 1595 | 36.48 | 5.81 x 10<sup>-10</sup> | 2.97 x 10<sup>-8</sup> |
| YMR207C/HFA1 | 77 | 16 | 2215 | 2103 | 4.57 | 6.16 x 10<sup>-10</sup> | 3.08 x 10<sup>-8</sup> |
| YOR371C/GPB1 | 79 | 18 | 347 | 323 | 4.09 | 1.42 x 10<sup>-8</sup> | 4.02 x 10<sup>-7</sup> |
| YML099C/ARG81 | 96 | 16 | 351 | 247 | 4.22 | 1.61 x 10<sup>-8</sup> | 4.43 x 10<sup>-7</sup> |
| YJL062W/LAS21 | 74 | 10 | 170 | 131 | 5.70 | 2.19 x 10<sup>-8</sup> | 5.77 x 10<sup>-7</sup> |
| YPL268W/PLC1 | 112 | 16 | 245 | 146 | 4.17 | 4.57 x 10<sup>-8</sup> | 1.05 x 10<sup>-6</sup> |
| YCR042C/TAF2 | 155 | 45 | 661 | 470 | 2.45 | 1.70 x 10<sup>-7</sup> | 2.90 x 10<sup>-6</sup> |
| YPL254W/HFI1 | 76 | 10 | 150 | 106 | 5.37 | 1.66 x 10<sup>-7</sup> | 2.90 x 10<sup>-6</sup> |
| YPR049C/ATG11 | 131 | 45 | 448 | 395 | 2.57 | 1.67 x 10<sup>-7</sup> | 2.90 x 10<sup>-6</sup> |
| YNL049C/SFB2 | 32 | 1 | 773 | 587 | 24.30 | 3.22 x 10<sup>-7</sup> | 4.88 x 10<sup>-6</sup> |
| YKL114C/APN1 | 44 | 3 | 120 | 87 | 10.63 | 9.80 x 10<sup>-7</sup> | 1.19 x 10<sup>-5</sup> |
| YIL068C/SEC6 | 80 | 5 | 271 | 111 | 6.55 | 1.36 x 10<sup>-6</sup> | 1.52 x 10<sup>-5</sup> |
| YGR198W/YPP1 | 79 | 25 | 295 | 282 | 3.02 | 1.97 x 10<sup>-6</sup> | 2.10 x 10<sup>-5</sup> |
| YDR375C/BCS1 | 67 | 4 | 107 | 54 | 8.45 | 2.20 x 10<sup>-6</sup> | 2.28 x 10<sup>-5</sup> |
| YKL017C/HCS1 | 78 | 13 | 219 | 143 | 3.92 | 3.56 x 10<sup>-6</sup> | 3.28 x 10<sup>-5</sup> |
| YDR235W/PRP42 | 61 | 7 | 177 | 109 | 5.37 | 5.34 x 10<sup>-6</sup> | 4.57 x 10<sup>-5</sup> |
| YDR180W/SCC2 | 173 | 72 | 523 | 431 | 1.98 | 6.47 x 10<sup>-6</sup> | 5.35 x 10<sup>-5</sup> |
| YMR167W/MLH1 | 94 | 26 | 286 | 209 | 2.64 | 2.47 x 10<sup>-5</sup> | 1.54 x 10<sup>-4</sup> |
| YKL197C/PEX1 | 132 | 58 | 415 | 351 | 1.92 | 1.55 x 10<sup>-4</sup> | 6.57 x 10<sup>-4</sup> |
| YMR078C/CTF18 | 73 | 26 | 295 | 260 | 2.47 | 1.63 x 10<sup>-4</sup> | 6.82 x 10<sup>-4</sup> |
| YGL095C/VPS45 | 67 | 14 | 202 | 122 | 2.89 | 3.72 x 10<sup>-4</sup> | 1.30 x 10<sup>-3</sup> |
| YMR094W/CTF13 | 54 | 21 | 152 | 158 | 2.67 | 4.44 x 10<sup>-4</sup> | 1.50 x 10<sup>-3</sup> |
| YNL132W/KRE33 | 75 | 4 | 187 | 52 | 5.21 | 5.13 x 10<sup>-4</sup> | 1.69 x 10<sup>-3</sup> |
| YDR103W/STE5 | 93 | 56 | 344 | 373 | 1.80 | 1.57 x 10<sup>-3</sup> | 4.16 x 10<sup>-3</sup> |
| YOR092W/ECM3 | 16 | 1 | 648 | 498 | 12.30 | 1.92 x 10<sup>-3</sup> | 4.86 x 10<sup>-3</sup> |
| YNR045W/PET494 | 61 | 20 | 177 | 136 | 2.34 | 2.15 x 10<sup>-3</sup> | 5.32 x 10<sup>-3</sup> |
| YJR107W/LIH1 | 19 | 1 | 102 | 61 | 11.36 | 2.40 x 10<sup>-3</sup> | 5.79 x 10<sup>-3</sup> |
| YLR397C/AFG2 | 102 | 27 | 288 | 155 | 2.03 | 2.56 x 10<sup>-3</sup> | 6.10 x 10<sup>-3</sup> |
| YGL082W/MIY1 | 41 | 8 | 132 | 84 | 3.26 | 2.60 x 10<sup>-3</sup> | 6.17 x 10<sup>-3</sup> |
| YOR091W/TMA46 | 40 | 6 | 103 | 46 | 2.98 | 0.0212 | 0.0351 |
| YDR456W/NHX1 | 87 | 13 | 187 | 59 | 2.11 | 0.0278 | 0.0443 |
| YDR508C/GNP1 | 3 | 0 | 959 | 797 | inf | 0.2562 | 0.3009 |
| YIL152W/VPR1 | 19 | 11 | 70 | 59 | 1.46 | 0.4184 | 0.4669 |

**Supplementary Table 10. Whole-gene tests for evidence of non-neutral protein evolution at thermotolerance loci.** Each row reports results from the McDonald-Kreitman test on sequences from strains of European populations of *S. cerevisiae* and *S. paradoxus* of the indicated top hit from barcoded RH-seq mapping of thermotolerance. D_s_, number of sites of synonymous nucleotide divergence between species; number of sites of D_n_, nonsynonymous nucleotide divergence between species; P_s_, number of sites of synonymous nucleotide polymorphisms within species; P_n_, number of sites of nonsynonymous nucleotide polymorphisms within species. NI, neutrality index. The sixth column reports the *p*-value from a Fisher’s exact test on D_s_, D_n_, P_s_, and P_n_, and the seventh column reports the adjusted *p*-value after applying the Benjamini-Hochberg correction for multiple hypothesis testing. All loci exhibited NI > 1, corresponding to a dearth of divergent amino acid changes relative to synonymous changes and polymorphisms.

