## Supplementary figures and images for "Barcoded reciprocal hemizygosity analysis via sequencing illuminates the complex genetic basis of yeast thermotolerance"

### Supplementary Figure 1

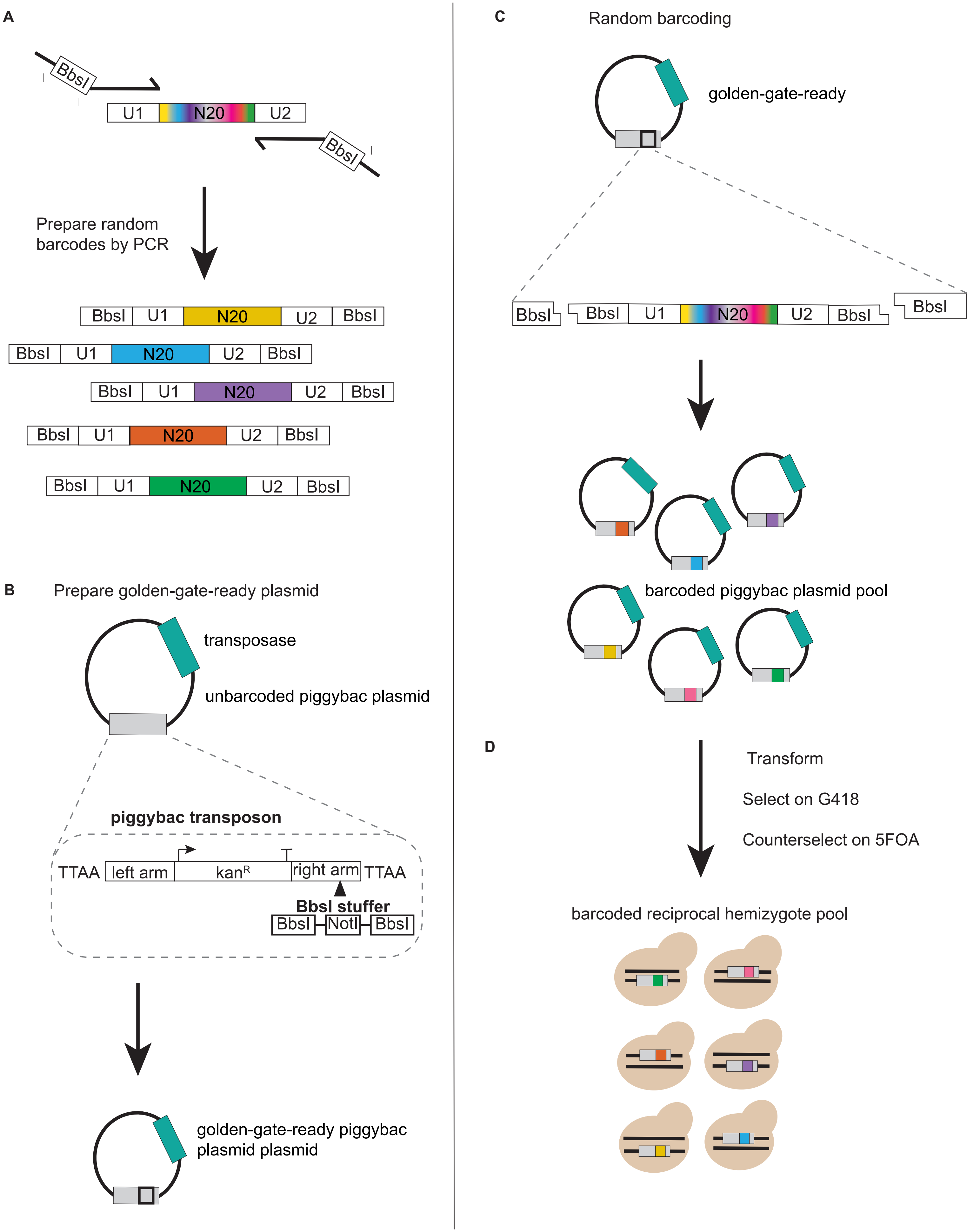

### Supplementary Figure 2

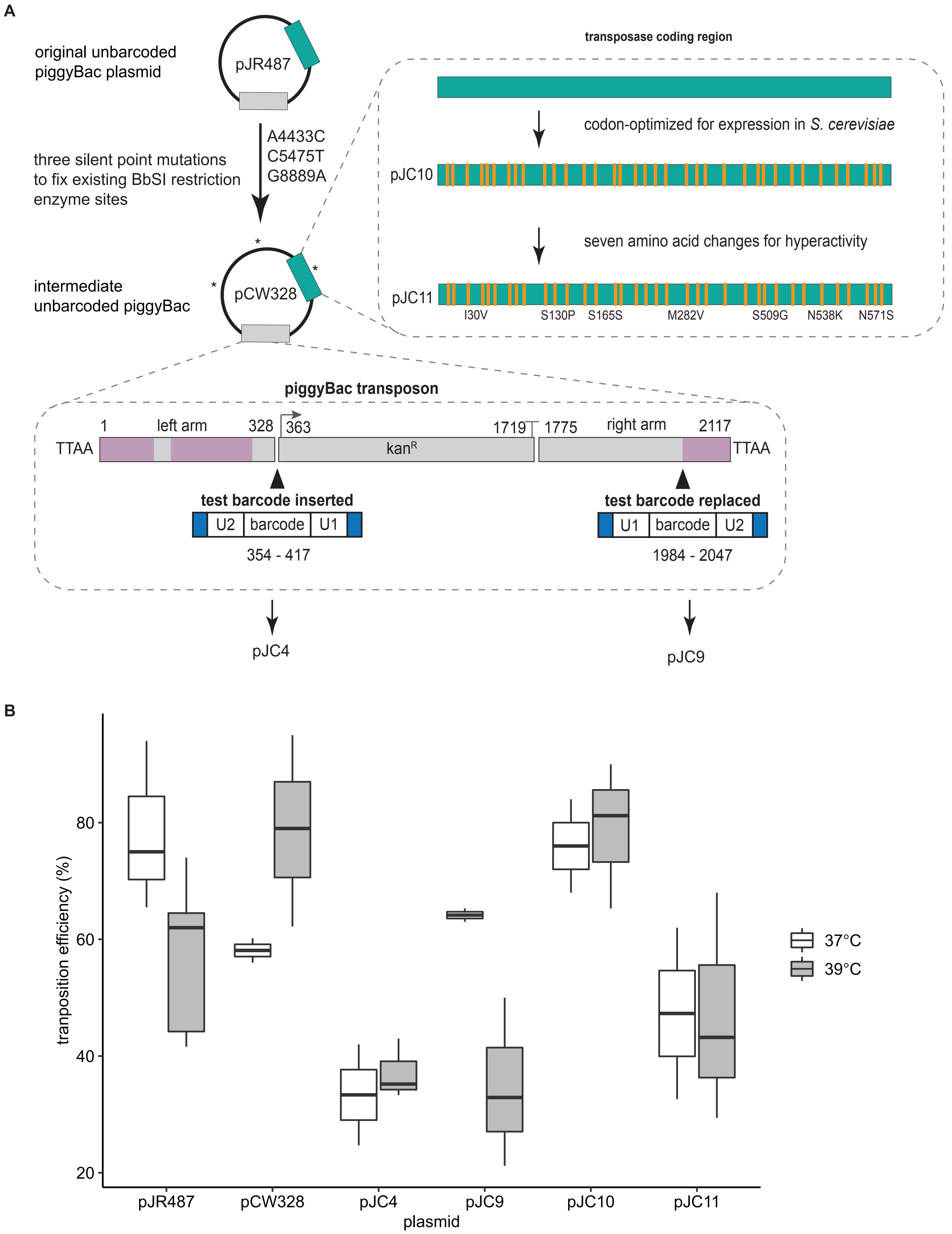

### Supplementary Figure 3

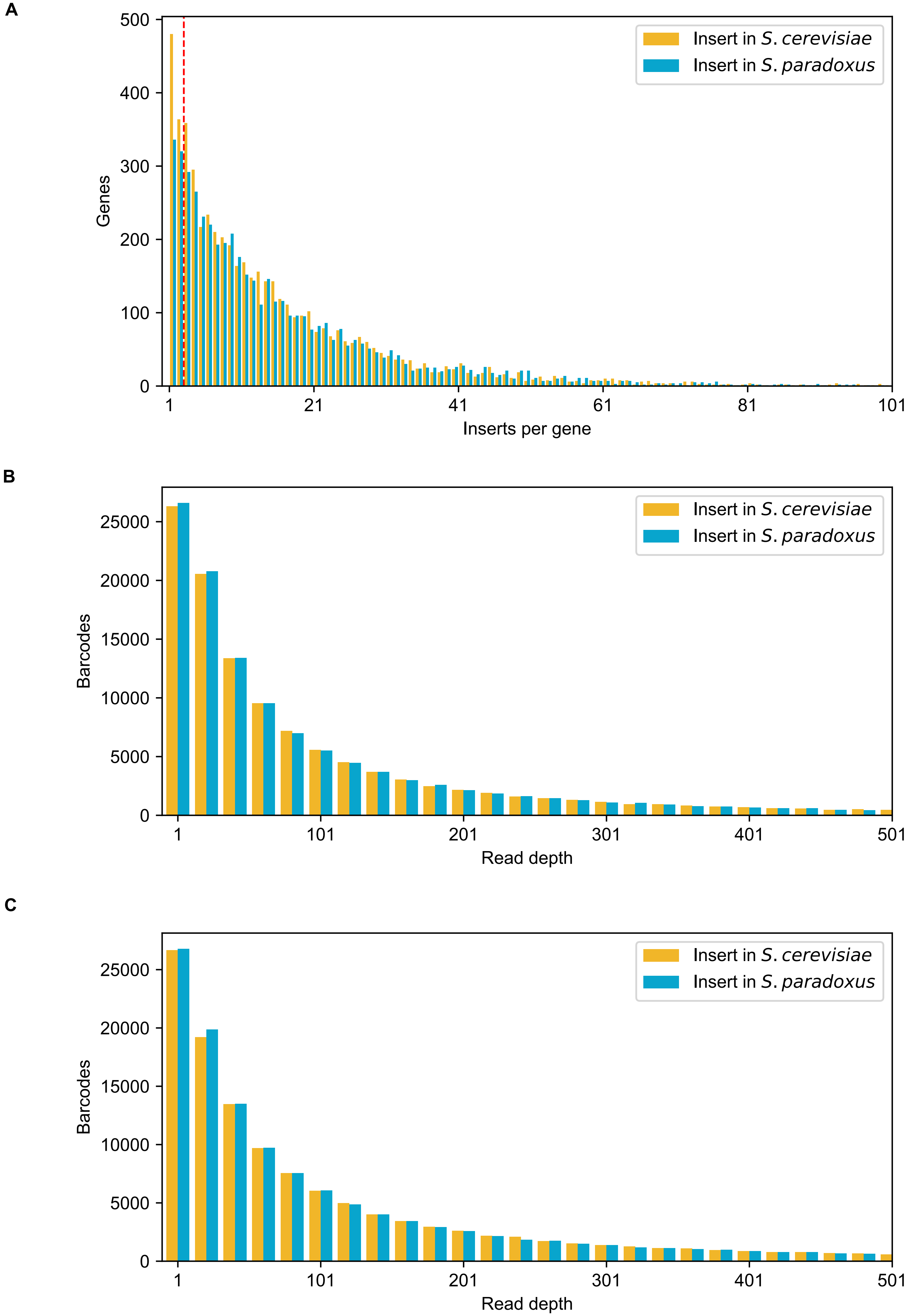

### Supplementary Figure 4

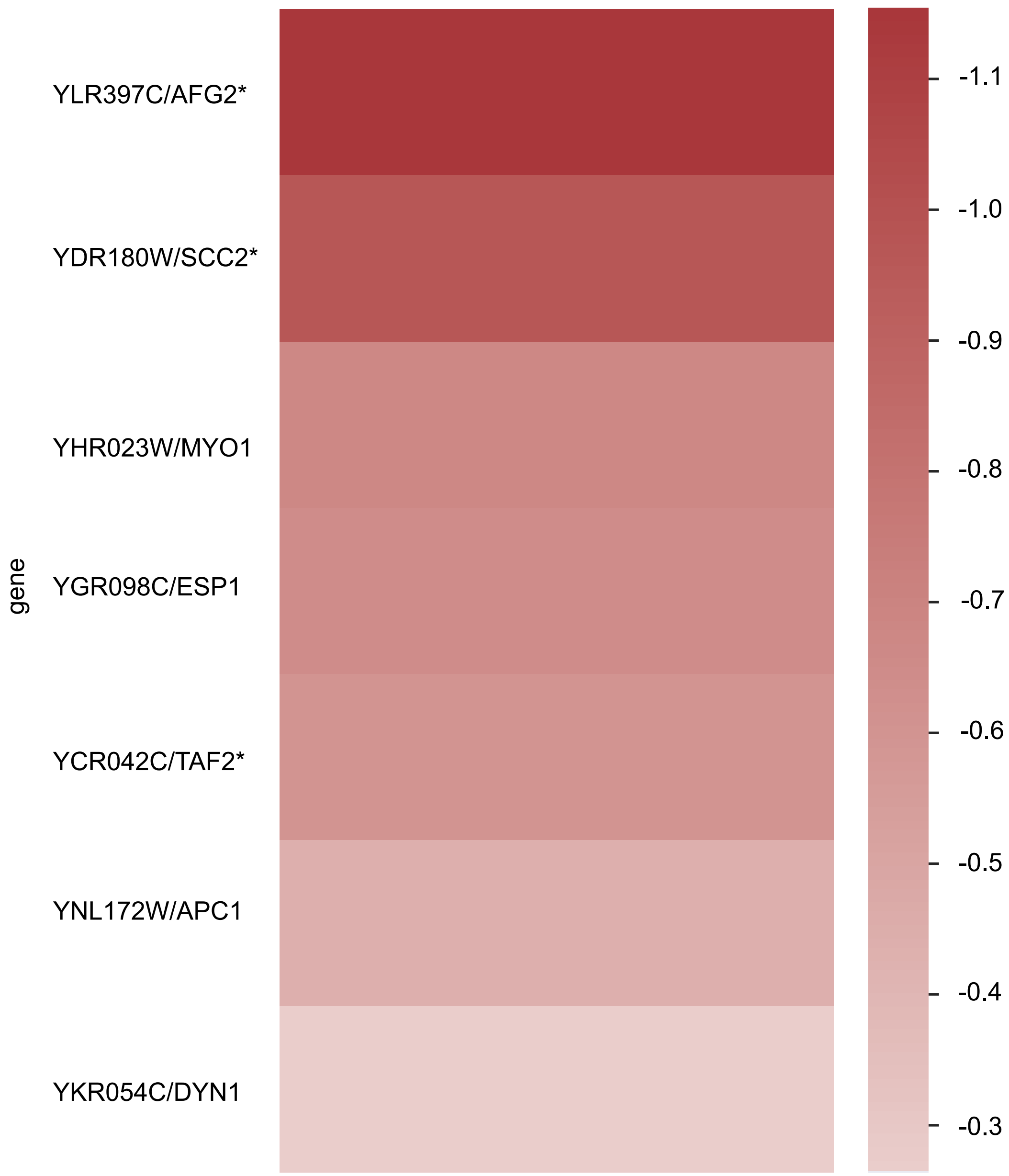

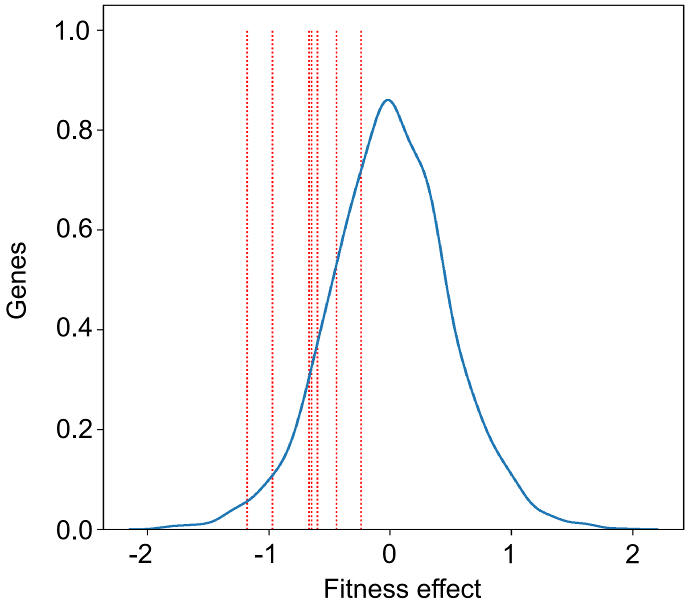

### Supplementary Figure 5

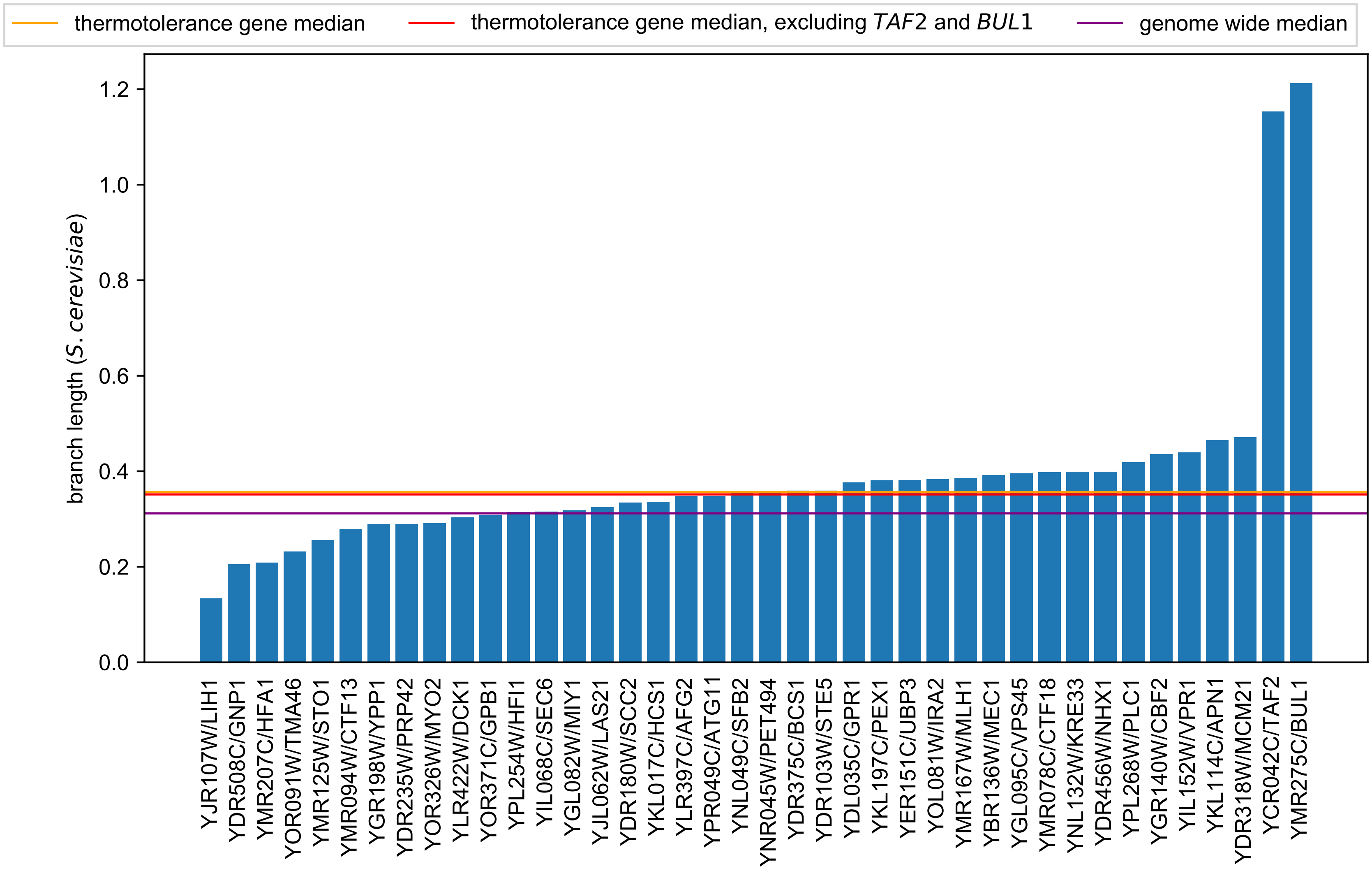
